# Neural representation of the decisional reference point in monkeys

**DOI:** 10.1101/2024.12.20.629801

**Authors:** Duc Nguyen, Erin L. Rich, Joni D. Wallis, Kenway Louie, Paul W. Glimcher

## Abstract

The decisional reference point serves as a hidden benchmark for evaluating options in decision-making. Despite extensive behavioral evidence for the existence of the reference point, its neural instantiation remains unclear. To identify reference point encoding at both the single-neuron and population levels, we analyzed neural activity from macaques performing a wealth accumulation task that dissociated objective reward values from the reference point. Across six frontal regions, we found that reference-related signals were broadly distributed at the single-neuron level. However, at the population level, only the ventral bank of the anterior cingulate cortex (vbACC) encoded the reference point significantly. In contrast, the dorsal bank of ACC and dorsolateral prefrontal cortex showed population-level encoding of reference-dependent subjective value signals. The temporal sequence of these signals and their known anatomical connectivity hints at a dedicated neural circuit for reference-dependence, with the vbACC potentially serving as a global source of reference point signal modulating activity in downstream value-encoding regions.

**SIGNIFICANCE:** All experiences are evaluated relative to an internal benchmark called the reference point. Despite its central role in all modern theories of decision-making, its neural basis has remained unclear. In this study, using dense single-neuron recordings from macaque frontal cortex, we found that only the ventral bank of the anterior cingulate cortex encoded the reference point consistently at the population level. We also found population-level reference-dependent value signals representing anticipated and earned rewards in the dorsal bank of the anterior cingulate cortex and the dorsolateral prefrontal cortex, respectively. Both were inversely modulated by the reference point as predicted by theory. These findings together with known anatomical connectivity support the existence of a dedicated neural circuit for reference-dependent valuation.

## INTRODUCTION

A key insight from modern behavioral decision-making is that people evaluate rewards and punishments not in absolute terms, but relative to an internal benchmark known as the ***reference point***. This idea distinguishes modern theories from classical ones, which assumed fixed preferences based on the final outcomes of one’s choices. Critically, the classical approach failed to capture the moment-by-moment fluctuations in the amount of reinforcement and anticipation we experience from a given reward in daily life as our environment changes. This limitation inspired Kahneman and Tversky to develop their landmark theory, Prospect Theory (PT), first formally introduced in 1979 (1). PT proposed that the subjective values which guide decision-making are inherently labile because they depend on how outcomes are compared to a context-dependent reference point. This framework successfully explained behaviors that classical theories could not (Fig. 1A) and revolutionized economic thought (1–3).

**Figure 1:**
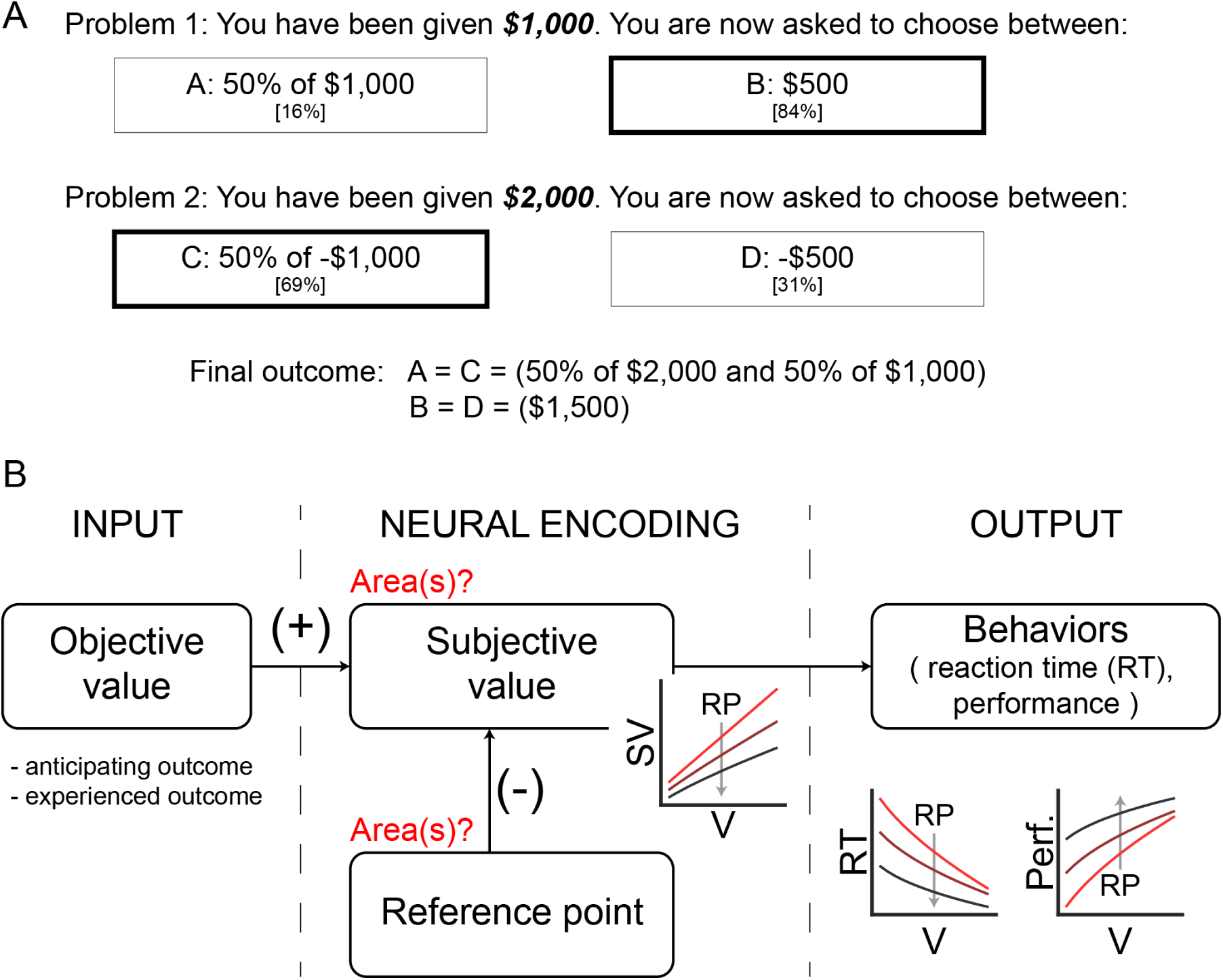
Demonstration of reference point and hypothetical subjective value computation following Kahneman and Tversky (1979). A. Example of reference-dependent choice behavior from Kahneman and Tversky 1979. Problem 2 is obtained from Problem 1 by moving $1,000 from all outcomes to the initial amount you are given. In terms of final outcomes, options A and C are equivalent, as are B and D. In Problem 1, however, a majority of people choose B (84%), whereas in Problem 2, a majority of people choose C (69%). Classical expected utility theories failed to explain this behavior because they were only influenced by final values of options. Prospect Theory, in contrast, proposed that subjective value is calculated as a function of both the initial endowment (here the *reference* point) and objective outcomes. B. Hypothetical subjective value (SV) computational circuit. In Prospect theory, people evaluate values (V) relative to their internal benchmark, called the reference point (RP). This personal RP exerts a negative influence on subjective value: higher RPs lower the perceived value of outcomes. Hence, the RP can modulate behaviors directly by itself or indirectly through modulating the perception of subjective values. We ask specifically if there exists brain area(s) that encode the RP on the population-average neural activity level.

In PT and related frameworks that descend from it, the reference point is an internal and dynamic benchmark constructed by the past and current environment. When this reference is low, even a small reward can rise to desirability, but when it is high, that same reward might lie well below the reference point and instead be perceived as undesirable. Decades of behavioral research have confirmed that decision-making is fundamentally reference-dependent (3–6). Inspired by these insights, neurobiological studies have revealed that the brain encodes reference-dependent value signals (7, 8), and that reward history shapes adaptive neural responses across multiple regions (9–14).

Despite extensive behavioral and neural research on reference-dependence, how the brain establishes or encodes the reference point remains unclear. In this study, we tested two hypotheses: (1) that single fronto-cortical neurons encode the reference point in their firing rates, and (2) that one or more regions show population-level encoding of the reference point, suggesting area-level specialization. That such an area may exist is compatible with Prospect Theory which proposes that the subjective value of an outcome is computed by subtracting the reference point from its objective value (Fig. 1B). This proposed calculation suggests that a homogeneous representation of the reference point could arise as the output of a brain region, which computes this signal and distributes it globally for value computation across the brain. A strong candidate for such a region is the anterior cingulate cortex (ACC), where reference-related signals have been observed during foraging tasks in multiple species (15–18).

To test these hypotheses, we reanalyzed neuronal recordings from Rich and Wallis (19). In this dataset, monkeys performed a wealth-accumulation task aimed at maximizing reward over a block of trials. The accumulated reward is, by definition, the reference point in Prospect Theory and related frameworks. Importantly, the design of this task allowed us to separate trial-by-trial reference point from the reward values on each trial.

Our findings revealed a critical double dissociation. At the single-neuron level, reference-related signals were broadly distributed across frontal regions without clear areal specialization. In contrast, when examined at the population-level, only the ventral bank of the ACC (vbACC) encoded the behaviorally specified reference point homogeneously across the neuronal population. In addition, we found a population-level reference-dependent cue valence encoding only in the dorsal bank of the ACC (dbACC), and a population-level reference-dependent outcome signal was observed only in the dorsolateral prefrontal cortex (dlPFC). In both dbACC and dlPFC, the value-related neural responses decreased as the reference point increased, consistent with predictions from Prospect Theory. This temporal and anatomical progression mirrors known projection pathways, supporting the existence of a dedicated frontal circuit for reference-dependent value computation in decision-making.

## RESULTS

### Wealth-accumulation task

To investigate neural encoding of the reference point, we reanalyzed data from previously published studies (19, 20). Prior analyses showed that multiple frontal brain regions in monkeys encode various decision-related variables but had not been specifically examined with respect to the instantaneous accumulated reward, which following Kahneman-Tversky (1), we defined as the reference point.

Two male *Macaca mulatta* subjects (age 6 and 10) were trained to perform an instructed response task over several hundred trials per session. Briefly after the start of each trial, a fixation point appeared, requiring the monkeys to fixate for 650ms. Fixation was followed by a visual cue, indicating whether the trial involved a potential gain or loss and prompted a joystick movement (left or right) to achieve the best possible outcome (either gaining or not losing reward). Depending on cue valence and the monkey’s response, feedback was provided by showing whether the accumulated amount had increased, decreased, or remained unchanged. The cumulative amount of reward subjects earned was represented by the length of the accumulated-reward bar that remained visible throughout the trial and served as the reference point for evaluating individual trial outcomes (Fig. 2A).

**Figure 2:**
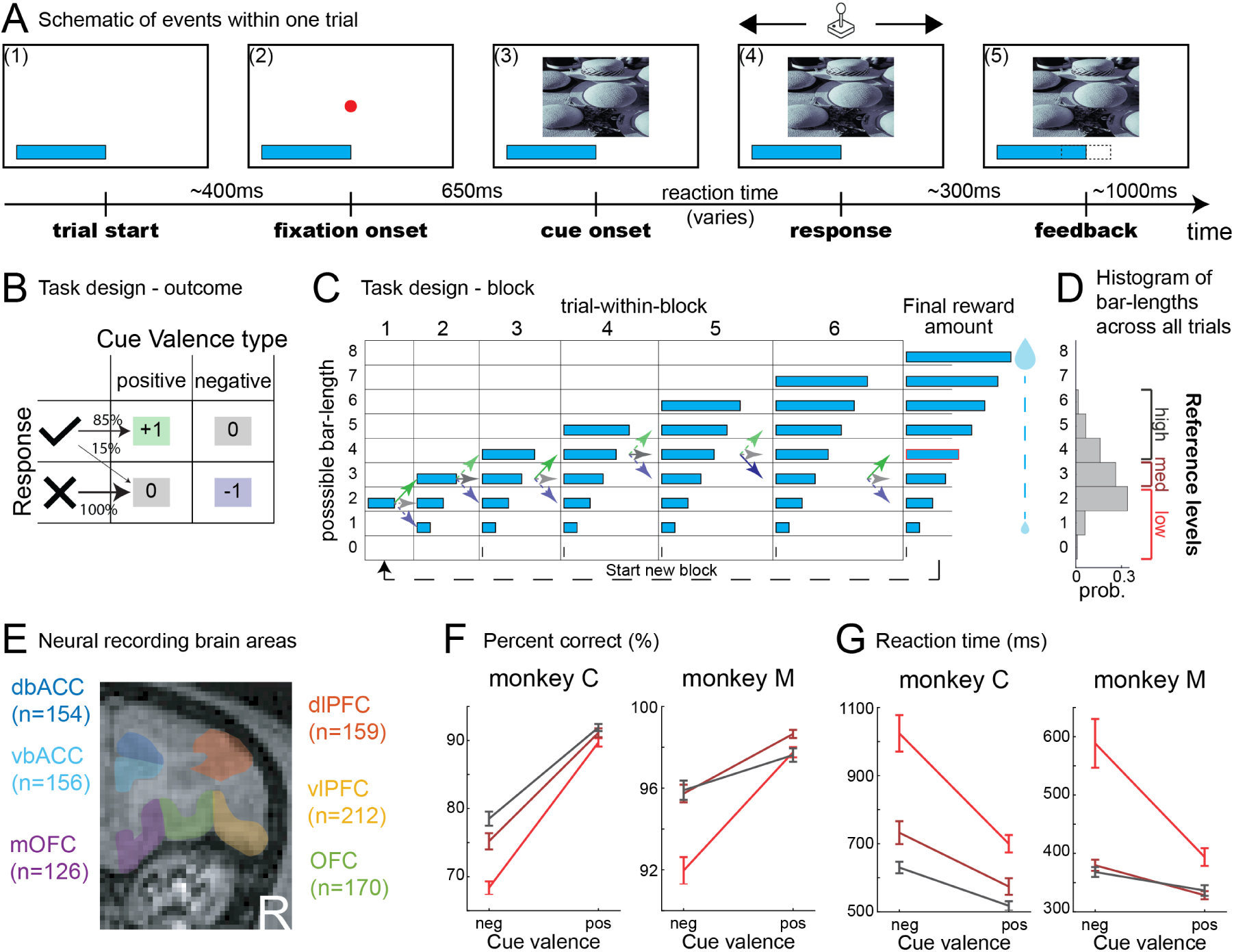
Instructed response task with accumulating reward. A. Schematic diagram of instructed response task. (1) The trial started after a random inter-trial-interval. The accumulated-reward bar remained visible throughout a block of 6 trials and only disappeared during reward delivery at the end of the block. Its length indicated the cumulative amount of reward subjects earned at that time point and served as a reference point against which individual trial outcomes could be evaluated. (2) Fixation point onset instructing subjects to fixate on it to start the task. (3) One of four familiar pictures were randomly chosen to appear on the screen. (4) Subjects responded by moving a joystick to left or right. (5) After a ∼300ms delay, subjects received feedback via change (or no change) of the accumulated-reward bar length, indicating whether they gained/ lost reward or nothing changed. B. Task design: Outcome depends on subjects’ responses and cue valence. A total of 4 visual cues were used as stimulus. For two cues of positive valence, a correct response (check) probabilistically increased the reward amount (top-left) and an error (X) resulted in no change (bottom-left). For the two of negative valence cues, a correct response probabilistically resulted in no change (top-right) and errors resulted in decreased reward amount (bottom-right). The bar length could not decrease below 0, so when the current bar length was 0, errors always produced no change. C. Task design: Subjects completed blocks of 6 trials before reward realization. Blocks always started at the accumulated-reward bar length of 2. Subjects proceeded through 6 trials before receiving the amount of reward accumulated by the end of the 6th trial. The volume of juice reward at the end of the block was proportional to the length of the bar. Arrows indicate potential reward amount earned after finishing each trial, smooth arrows showed an example of one block. D. Distribution of lengths of accumulated-reward bar at the beginning of each trial. We clustered trials into low, medium and high reference level trials based on the initial bar-length of that one. E. Schematic diagram of recorded regions of the brain. Shown on the coronal section of the right hemisphere. In subject M, dbACC, vbACC, mOFC, and OFC neurons were recorded in the left hemisphere, and dlPFC, vlPFC, and OFC neurons were recorded on the right. In subject C, dbACC, vbACC, mOFC, and OFC neurons were recorded in the right hemisphere and dlPFC, vlPFC, and OFC neurons were recorded on the left. F. Percent correct of subjects separated by trials of different reference levels and cue valence. For statistics, see Tables 1-2. G. Reaction time of subjects separated by trials of different reference levels and cue valence. For statistics, see Tables 1-2. Error bars represent mean ± SEM.

**Table 1:** Fixed effects coefficients on behavioral performance.

A. Reaction time
| Regressor | Coefficient (SE) | p-value |
| --- | --- | --- |
| *intercept* | 1002.50 (108.33) | $2.27 \times 10^{-20}$ |
| Expected-reward bar length /<br>Reference point | -123.87 (6.05) | $8.63 \times 10^{-93}$ |
| Valence (positive-cue indicator) | -382.99 (25.10) | $2.00 \times 10^{-52}$ |
| Reference point x Valence | 78.685 (8.20) | $9.04 \times 10^{-22}$ |

B. Percent correct
| Regressor | Coefficient (SE) | p-value |
| --- | --- | --- |
| *intercept* | 1.24 (0.6128) | 0.0428 |
| Expected-reward bar length /<br>Reference point | 0.23 (0.0197) | $3.66 \times 10^{-32}$ |
| Valence (positive-cue indicator) | 1.66 (0.0973) | $2.93 \times 10^{-65}$ |
| Reference point x Valence | -0.16 (0.0335) | $2.43 \times 10^{-6}$ |

**Table 2:** Extended regression models on behavioral performance.

| Regressor | Coefficient (SE) | p-value |
| --- | --- | --- |
| *intercept* | 926.52 (94.37) | $1.02 \times 10^{-22}$ |
| Expected-reward bar length /<br>Reference point | -83.75 (6.73) | $1.81 \times 10^{-35}$ |
| Valence (positive-cue indicator) | -368.81 (25.40) | $1.25 \times 10^{-47}$ |
| Reference point x Valence | 75.50 (7.83) | $5.79 \times 10^{-22}$ |
| Direction (left-response indicator) | 52.06 (9.35) | $2.59 \times 10^{-8}$ |
| Trial number within a block | -8.31 (3.65) | 0.0228 |
| Trial number within session | 0.24 (0.02) | $9.62 \times 10^{-23}$ |
| Previous-trial reaction time | 0.07 (0.0048) | $2.15 \times 10^{-53}$ |
| Previous-trial cue valence | -11.69 (16.49) | 0.4786 |
| Previous-trial cue direction | 9.75 (9.36) | 0.2975 |
| Previous-trial response (correct-<br>response indicator) | -187.7 (19.84) | $3.33 \times 10^{-21}$ |
| Previous-trial reward | -11.93 (14.89) | 0.4232 |

**B. Percent correct**
| Regressor | Coefficient (SE) | p-value |
| --- | --- | --- |
| *intercept* | 1.25 (0.6103) | 0.0398 |
| Expected-reward bar length /<br>Reference point | 0.20 (0.0259) | $3.23 \times 10^{-15}$ |
| Valence (positive-cue indicator) | 1.67 (0.1120) | $2.33 \times 10^{-50}$ |
| Reference point x Valence | -0.18 (0.0362) | $7.65 \times 10^{-7}$ |
| Direction (left-response indicator) | 0.48 (0.0421) | $1.864 \times 10^{-29}$ |
| Trial number within a block | 0.01 (0.0155) | 0.4393 |
| Trial number within session | -0.0005 (0.0001) | $2.66 \times 10^{-6}$ |
| Previous-trial reaction time | -0.00006 (0.00002) | $4.56 \times 10^{-5}$ |
| Previous-trial cue valence | -0.37 (0.0705) | $1.49 \times 10^{-7}$ |
| Previous-trial cue direction | 0.08 (0.0417) | 0.0422 |
| Previous-trial response (correct-<br>response indicator) | 0.24 (0.0766) | 0.0014 |
| Previous-trial reward | 0.09 (0.0647) | 0.1791 |

The task used four trial-type cues in a 2×2 design, differing by correct response direction (left/ right) and valence (positive/ negative). Positive cues increased the reward if the response was correct and gave no reward if incorrect. Negative cues gave no reward if correct and reduced the reward if incorrect (Fig. 2B). Subjects completed six sequential trials per block before receiving their juice reward, an amount proportional to the length of the accumulated-reward bar displayed at the end of the sixth trial (Fig. 2C).

In this study, we explicitly tested two neural hypotheses that are associated with two core features of PT: (1) that the reference point is represented discretely in neural firing rates; and (2) that the neural representations of both anticipated and received outcomes are modulated by the current reference point value, reflecting the observed reference-dependent behaviors. To analyze the specific influence of the reference point on neuronal firing rates, we performed parametric linear regression analyses using the length of the accumulated-reward bar, representing the current reference point level, as a regressor. Following previous studies (e.g. (21)), we implicitly assumed a linear utility function which allows us to estimate the impact of the reference point on neuronal firing rates. While our experimental design did not allow a direct estimate of the curvature of the utility function, any non-linear impacts of the reference point would be absorbed by our regression constant across all conditions, allowing us to accurately detect reference effects within our linear constraints. For visualization, we further relaxed this linear constraint by non-parametrically grouping trials into three discrete reference levels: low, medium and high (Fig. 2D).

We recorded single neuronal activities from six frontal cortical areas. Of these, only neurons that had sufficient activity for statistical analyses and clearly identified anatomical location were used in our analysis (Fig. 2E and Methods).

#### I. Reference-dependent behaviors

Previous studies have shown that animals’ behavioral performance improves with both the magnitude of potential rewards (22, 23), equivalent to *cue valence* in this study, and the current motivational context, or the *reference point* (24, 25). However, theories also predict that higher reference point can impair performance by reducing the subjective value of rewards (1). To determine whether the behaviorally specified reference point influences reward-related behavior, we examined both *trial-by-trial reaction time* and *percent correct* as a function of the *reference point* using mixed-effects regression models (see Methods). Our results aligned with theoretical predictions (Table 1; Figs. 2F-G): higher reference points and higher potential rewards were associated with faster reaction times and higher percentage correct. Also as predicted, their interaction was associated with slower reaction times and lower percent correct rates.

To control for potentially confounding variables such as history (performance and outcome on previous trials) or the trial’s position within the block (counting down to reward delivery), we ran additional models including these factors (see Methods). The reference point effect remained robust under our most stringent controls (Table 2), confirming that behavioral performance in these animals is, as expected, reference-dependent.

#### II. Neural representation of the reference point

To examine the effect of the reference point itself or reference-dependent reward signals on neural activity, we focused on three keys epoch: *trial start*, *cue onset*, and *feedback*. Each successive epoch introduced additional decision-related information and reflected the dynamic evolution of the decision-making process.

To isolate the reference point signal, we first focused on the *trial start* epoch, when the accumulated-reward bar length (the current reference point) was the only available information (Fig. 3A). In this epoch, subjects had no knowledge of the upcoming trial type or the outcome, minimizing task-related confounds.

**Figure 3:**
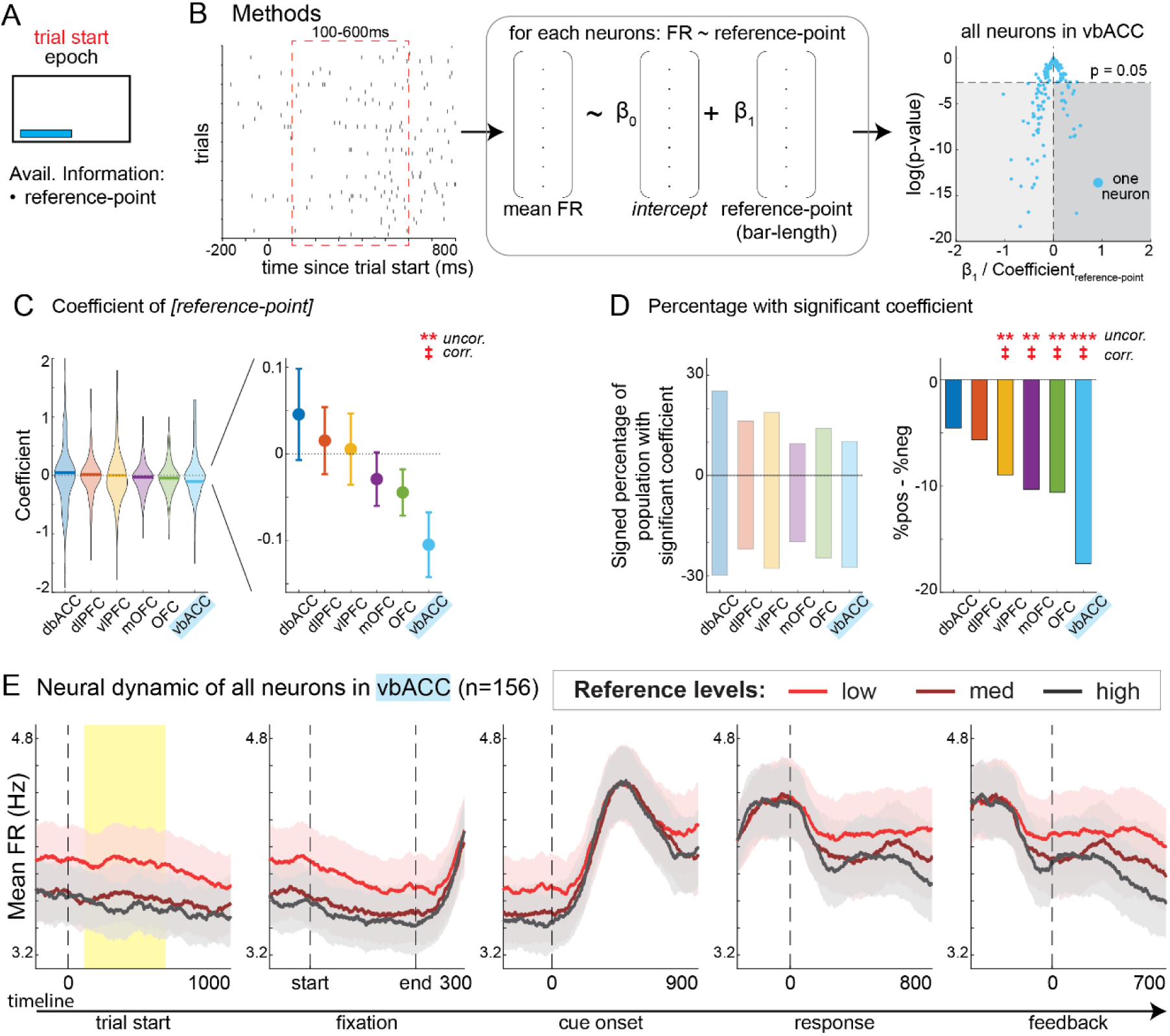
Population level encoding of the reference point in the ventral bank of anterior cingulate cortex (vbACC) during trial start epoch. A. Schematic of trial start epoch. The only information available to subjects is the current amount of accumulated reward, which acts as a reference point. Therefore, we fitted a regression model with bar-length (i.e. reference point). B. Schematic of analysis plan. (Left) Time window for analysis. (Middle) For each neuron, we averaged spike count in this window to fit regression models. (Right) From this, we obtained a coefficient value (β) and its p-value for each neuron for each regressor and use this for population analysis. We implemented two main tests: (1) t-test on population-average coefficients; and (2) χ^2^-test on percentage-of-positive vs. negative coefficients within significant populations. C. Distribution (left) and mean ± SEM (right) for all coefficient values of reference point (*bar-length*) of neurons grouped by brain area. P-value of one-sample t-test for vbACC: 0.0056. For all statistics, see Table S1. D. The percentage of population with significantly positive and negative coefficient values for reference point (*bar-length*) per brain area (left) and the difference between these two numbers (right). P-value of χ^2^test for positive versus negative: vlPFC: 0.0069; mOFC: 0.0025; OFC: 0.0017; vbACC: 6.66×10^−7^. For all statistics, see Table S1. E. Average firing rate of all neurons in vbACC across all events of a trial. Line plots indicate mean and shaded areas indicate SEM. Error bars represent mean ± SEM. *p ≤ 0.05, **p ≤ 0.01, ***p ≤0.001, n.s. p > 0.05. ‡p ≤ 0.0083 (Bonferroni-corrected for 6 brain areas).

##### a. Single-neuron level analysis across frontal regions

We conducted regression analyses on trial-by-trial firing rates to assess whether individual neurons encoded the reference point during this epoch (Fig. 3B; see Methods). Because no other task variables were present at this point, this analysis specifically targeted encoding of the reference point itself.

We found neurons with both positive and negative tuning to the reference point; increasing or decreasing firing rates as the reference point increased. This heterogeneous response pattern was observed across all six frontal cortical regions (Fig. 3C-left). We further quantified the proportion of neurons in each area showing significant positive or negative coefficients (Fig. 3D-left). While many neurons across regions encoded the reference point, this signal was broadly distributed and mixed across areas. These findings suggest that reference-related signals are widespread at the level of individual neurons, without a clear unique anatomical specialization.

To examine population structure, we performed two complementary analyses, principal component analyses (PCA) and demixed PCA (dPCA) (see Methods). Consistent with the single-neuron results, we found that activity in all regions formed low-dimensional manifolds that differentiate between reference levels, mainly along the first principal component (Fig. S1, left and middle columns). We then performed dPCA to separate reference encoding from temporal dynamics (26), we again observed distinct trajectories for each reference-level, confirming the reference-related signals in these regions (Fig S1, right column).

##### b. At population level, only the ventral bank of the anterior cingulate cortex (vbACC) encoded the reference point significantly

We next tested whether any brain region exhibited a population-level encoding of the reference point, a representation that could serve as a global signal for downstream value computations. To perform this test, for each area, we conducted two population-level analyses. First, we tested whether the average coefficient-of-reference-point values across neurons differed from zero. Among these areas, only the vbACC significantly deviated from zero (t-test, p = 0.0056), indicating consistent reference point encoding at the population level only in this area (Fig. 3C-right). Second, we examined the proportion of neurons with significant coefficients and found a strong bias toward negative encoding in the vbACC (χ^2^, p < 0.001), further supporting a homogenous representation in this region (Fig. 3D-right). Both conclusions held under uncorrected (p < 0.05) and Bonferroni-corrected thresholds (for 6 area comparisons, p < 0.0083).

These findings were robust across four different measures of firing rate (Fig. S2A; see Methods). Among all recorded regions using all of our four analytic approaches, only vbACC consistently showed population-level homogeneity, as evidenced by a biased distribution of regression coefficients (Fig. S2B) and a skewed proportion of neurons with significant reference point tuning (Fig. S2C).

Since monkeys were motivated to accumulate rewards throughout block, the reference point was correlated with block position (ρ = 0.48, p < 0.001), raising concerns about possible confounds. To address this, we first plotted population-average activity across reference-level and position-in-block combinations for each brain region. The vbACC activity mainly reflected reference levels (Fig. S3).

Next, we conducted a series of additional control analyses. First, to confirm this population-level result was not driven by differences across subjects or sessions, we performed a mixed-effects regression controlling for subject and session identity (see Methods). The vbACC again emerged as the only region with a significant and consistent effect (Figs. S4A-B).

To further control for other confounds, including the collinearity between reference point and the position of an individual trial in block or the drifting performance over time or influences from previous trials, we employed a hierarchical regression model. We first regressed out the influence of potential confounding factors, then modeled the residual variance using the reference point (see Methods). The main finding, that the reference point is encoded at the population-level in vbACC, remained significant under uncorrected thresholds for both tests, though some results did not survive Bonferroni correction (Figs. S4C-D). We then combined both controls, confounds and subject-level differences, and again found that the effect persisted under uncorrected thresholds but fell short of corrected significance (Figs. S4E-F). These results support the robustness of reference point encoding in the vbACC at the population level.

While we focused on the objective accumulated reward as the reference point, it is possible that monkeys used a subjective internal benchmark that evolves try-by-trial. To explore this possibility, we tested four alternative models of subjective reference point dynamics. While some alternative models explained behavior or single-neuron activity better than the objective accumulated reward model, all converged on the same key conclusion: only the vbACC consistently encodes the reference point at population level (see Supplemental Note 1).

Finally, to visualize this population-level effect, we grouped trials by reference point levels and plotted average neuronal firing rates over time (Fig. 3E). These plots show that the influence of the reference point persists throughout the trial, as confirmed by repeated-measures ANOVA (Fig. S5A-B). Univariate regression analyses further demonstrated that this homogeneous encoding is stable across most epochs, except during the phasic transient elicited by cue onset (Fig. S5C). Together, these results provide strong evidence that the vbACC carries a dedicated, homogeneous representation of the reference point at the population level.

#### III. Neural representation of the reference-dependent cue valence

We next focused on the *cue onset* epoch, when subjects received new information in the form of a visual cue indicating a potential gain or loss. Importantly, the accumulated-reward bar remained visible during this period, allowing us to assess how it modulated the perception of potential future reward, equivalent to a reference-dependent *cue valence* signal (Fig. 4A).

**Figure 4:**
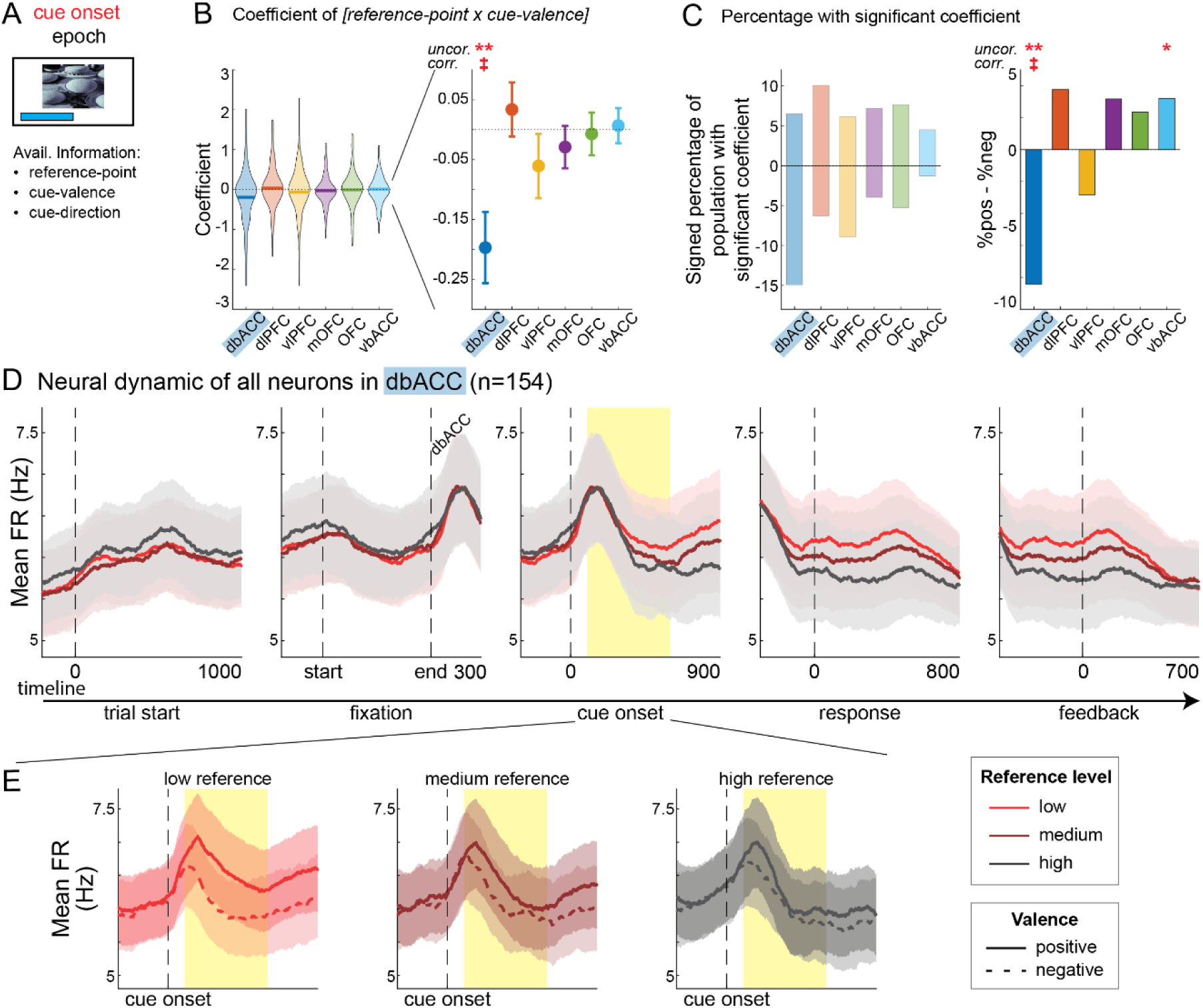
Population level encoding of the reference-dependent cue valence in the dorsal bank of anterior cingulate cortex (dbACC) during cue-onset epoch. A. Schematic of cue-onset epoch. Available information to subjects is current amount of reward that acts as a reference point, and the 2 properties of cue: cue valence and cue direction. For this epoch, we fitted a multivariable regression model that consists of the above 3 parameters and their interaction terms. B. Distribution (left) and mean ± SEM (right) for all coefficient values of reference-dependent cue valence signal (interaction term [*reference-point × cue-valence*]) of neurons grouped by brain area. P-value of one-sample t-test for dbACC: 0.0011. For all statistics, see Table S2. C. The percentage of population with significantly positive and negative coefficient values for reference-dependent cue valence signal (interaction term [*reference-point × cue-valence*]) per brain area (left) and the difference between these two numbers (right). P-value of χ^2^ test: dbACC: 0.0014; vbACC: 0.0184. For all statistics, see Table S2. D. Average firing rate of all neurons in dbACC across all events of a trial, separated by reference levels. Line plots indicate mean and shaded areas indicate SEM. Same in panel E. E. Average neural dynamics within cue-onset epoch of all neurons in dbACC, separated by reference levels and cue valence type. Error bars represent mean ± SEM. *p ≤ 0.05, **p ≤ 0.01, ***p ≤0.001, n.s. p > 0.05. ‡p ≤ 0.0083 (Bonferroni-corrected for 6 brain areas).

##### a. Single-neuron level analysis across frontal regions

For each neuron, we performed regression analyses including an interaction term between reference point and cue valence, allowing us to examine how subjective evaluation of potential reward was modulated by current reference point. Theoretical models predict a negative modulation of cue valence.

As with the reference point itself, we found neurons exhibiting both positive and negative tuning for this interaction term across all six frontal regions (Fig. 4B-left). We identified neurons with statistically significant coefficients and plotted the proportions of positive and negative tuning (Fig. 4C-left). Critically, all regions showed a mix of tuning for the reference-dependent cue valence encoding at the single-neuron level.

To examine the population structure of the representation of these values, we applied PCA and dPCA. Consistent with the single-neuron results, neural activity in all regions reflected a linear combination of reference point level and cue valence, primarily along the first and second principal components. dPCA revealed distinct trajectories for each reference-level and cue-valence condition (Fig. S6). Together, these findings indicate that reference-dependent cue-valence encoding is broadly distributed across the frontal cortex at single neuron level.

##### b. At population level, only the dorsal bank of the anterior cingulate cortex (dbACC) encoded a reference-dependent cue valence signal significantly

To test for a more selective encoding at the population level, we applied the same two analyses used in previous epoch. Among all brain regions, only the dbACC showed homogeneous, population-level, encoding of the reference-dependent cue valence signal. First, the average regression coefficient for this interaction term significantly differed from zero (t-test, p = 0.0011), indicating a consistent modulation of cue-valence-related activity by the reference point (Fig. 4B-right). Second, among neurons with significant coefficients, there was a strong bias toward negative tuning (χ^2^, p = 0.0014), consistent with theoretical predictions and further supporting population-level homogeneity (Fig. 4C-right). These findings were robust across different firing rate measures (Figs. S7A-B).

To confirm this effect, we tested for subject and session variability using a mixed-effects model. Most effects remained significant, although a few did not (Figs. S7C–D), possibly due to differences between parametric and non-parametric test assumptions.

We then ruled out confounding variables by first regressing out the same set of covariates employed in the previous epoch, then reapplying the main regression model to the residual firing rates (see Methods). The reference-dependent cue valence signal in dbACC remained significant, even under Bonferroni correction (Figs. S7E-F). When combining both controls, the effect remained robust under the strictest test (Figs. S7G-H). These results provide strong support for a homogeneous population-level encoding of reference-dependent cue valence in the dbACC.

To visualize the population-level effect, we first plotted the average firing rates of dbACC neurons across epochs grouped by reference levels (Fig. 4D). Neural responses differed across reference levels only during the cue onset epoch, but not before or after that epoch. We further separated the neuronal activities within the cue onset epoch by cue valence (Fig. 4E). We observed the largest difference in population responses to positive versus negative cues at low reference level. Consistent with reference-dependence theories, this valence difference diminished as reference point increased. Together, these findings offer evidence that the dbACC encodes a theory-consistent, reference-dependent cue valence signal at population level.

#### IV. Neural representation of the reference-dependent outcome

Lastly, we examined reference-related signals during the feedback epoch, when subjects received outcome information indicating whether their cumulative reward increased, decreased, or remained unchanged based on their action (Fig. 5A). This allowed us to test how the reference point modulated subjective evaluation of actual outcomes.

**Figure 5:**
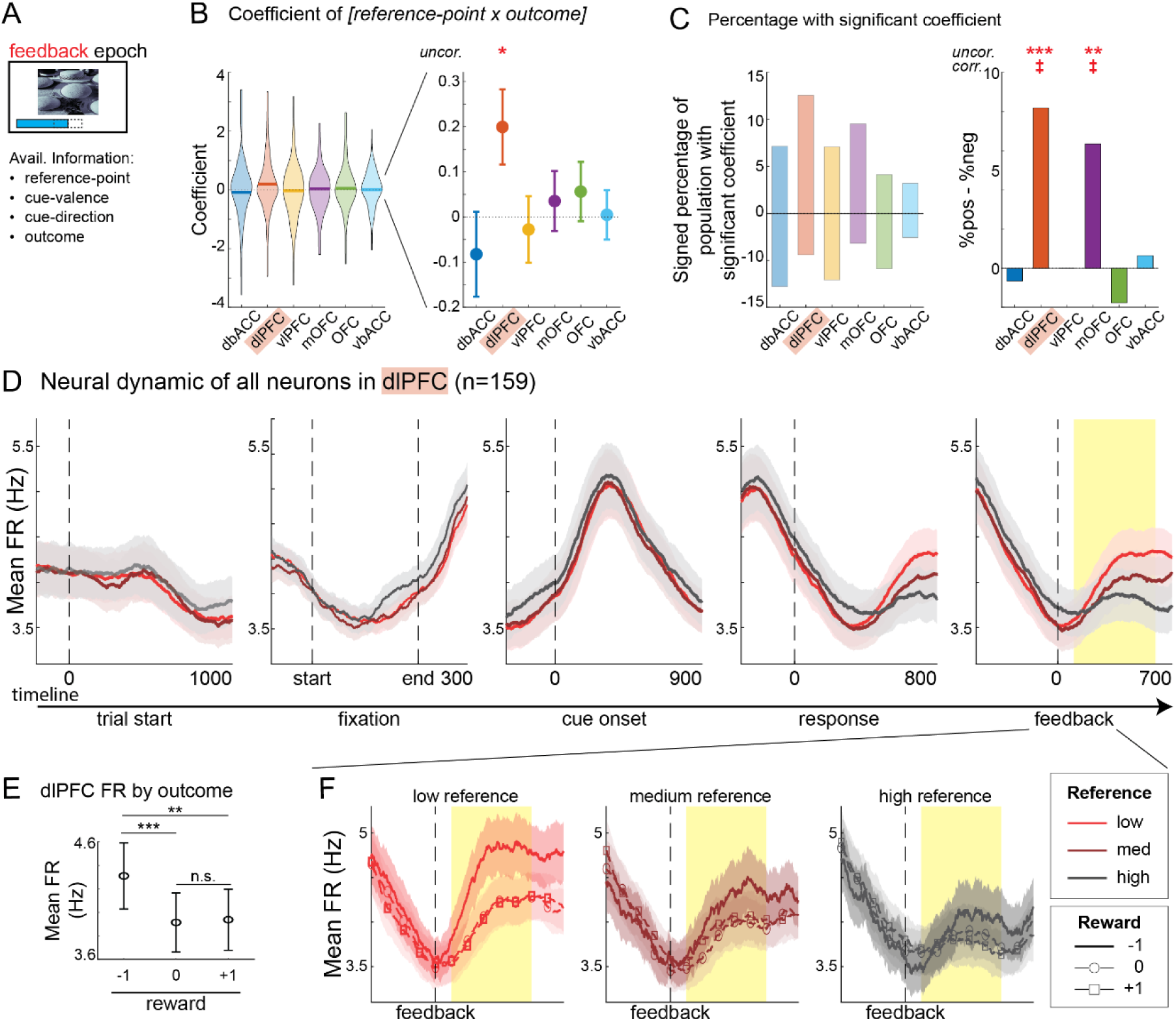
Population level encoding of the reference-dependent outcome signal in dorsolateral prefrontal cortex (dlPFC) during feedback epoch. A. Schematic of feedback epoch. Available information to subjects includes the current amount of reward that acts as a reference point, the 2 properties of the cue (cue valence and cue direction), and the outcome (change in earned reward). For this epoch, we fitted a multivariable regression model that consists of the above 4 parameters and their interaction terms. B. Distribution (left) and mean ± SEM (right) for all coefficient values of reference-dependent outcome signal (interaction term [*reference-point × outcome*]) of neurons grouped by brain area. P-value of one-sample t-test for dlPFC: 0.0176. For all statistics, see Table S3. C. The percentage of population with significantly positive and negative coefficient values for reference-dependent outcome signal (interaction term [*reference-point × outcome*]) per brain area (left) and the difference between these two numbers (right). P-value of χ^2^test: dlPFC: 4.03×10^−4^; mOFC: 0.0047. For all statistics, see Table S3. D. Average firing rate of all neurons in dlPFC across all events of a trial, separated by reference levels. Lines indicate mean and shaded areas indicate SEM. Same in panel F. E. The average firing rate of all neurons in dlPFC, showing it responded specifically to lost reward. P-values for (−1) vs. (0): 2.60×10^−5^; (−1) vs. (+1): 0.0011. F. Average neural dynamics within the feedback epoch of all neurons in dlPFC, separated by reference levels and earned reward amount. Error bars represent mean ± SEM. *p ≤ 0.05, **p ≤ 0.01, ***p ≤0.001, n.s. p > 0.05. ‡p ≤ 0.0083 (Bonferroni-corrected for 6 brain areas).

##### a. Single-neuron level analysis across frontal regions

For each single neuron, we used regression models to quantify the interaction between reference point and outcome (see Methods), capturing the reference-dependent modulation of outcome signals.

As with previous epochs, we observed neurons with both positive and negative tuning to the reference-dependent outcome signal in all six frontal regions (Fig. 5B-left). We then identified neurons with statistically significant coefficients and plotted the proportions of selectivity (Fig. 5C-left). PCA confirmed these findings, showing that neural activity reflected a linear combination of reference point and outcome levels, especially along the first two principal components. dPCA further revealed distinct trajectories for each reference-outcome condition (Fig. S8). Overall, no brain region showed a clear specialization, following a familiar pattern: reference-related modulation of outcome encoding is widespread and heterogeneous across the frontal cortex at the single-neuron level.

##### b. At population level, only the dorsolateral prefrontal cortex (dlPFC) encoded a reference-dependent outcome signal significantly

To identify region-specific encoding, we tested for population-level effects. Among all brain regions, only the dlPFC showed consistent, population-level, encoding of the reference-dependent outcome signal. First, the average regression coefficient for the interaction term (reference point × outcome) significantly differed from zero (t-test, p = 0.0176), indicating a consistent impact of the reference point on outcome encoding (Fig. 5B-right). Second, among neurons with significant coefficients, a greater proportion showed positive tuning, supporting a biased, homogeneous representation in this population (χ^2^, p < 0.001; Fig. 5C-right). These findings were robust across all firing rate metrics used (Figs. S9A-B).

To assess robustness, we first tested for subject- and session-level effects using a mixed-effects model. Most, though not all, effects remained significant (Figs. S9C–D). Next, we addressed other confounds by regressing out relevant covariates and applying the main model to the residual firing rates (see Methods). The reference-dependent outcome signal in dlPFC remained significant, even under Bonferroni correction (Figs. S9E–F). When combining both subject-level and covariate controls, the effect persisted under this strictest test (Figs. S9G–H). These findings support a homogeneous influence of the reference point on outcome encoding in the dlPFC at population level.

Finally, population-averaged firing rates showed that dlPFC activity reflected reference levels specifically during the feedback epoch (Fig. 5D). dlPFC responses were especially strong to negative outcomes (loss of earned reward), but, consistent with reference-dependence theories, this response diminished as the reference level increased (Figs. 5E-F).

## DISCUSSION

### Summary

While extensive behavioral evidence has shown that the reference point shapes both human and animal decision-making, and prior electrophysiological and imaging studies have demonstrated reference-dependent neural activity (7, 8, 27–29), the reference point itself has not yet been explicitly localized in the brain. At the single neuron level, we found reference-related signals distributed across multiple frontal brain regions, indicating broad but anatomically nonspecific encoding. These findings agree with prior studies that used similar task structures and recorded neurons from other brain regions, from subcortical to cortical areas in monkeys (8, 30–32).

In contrast to this broad distribution, at the population level, we found evidence for a much more selective level of representation. We found significant evidence for a distinctive population-level representation of the reference point only in the vbACC. Albeit at a slightly lower significance level, we also found evidence for population-level signals encoding a reference-dependent cue valence signal in the dbACC and a reference-dependent outcome signal in the dlPFC.

These results revealed a temporal and anatomical progression: the vbACC encoded the reference point early in the trial, followed by its influence on cue evaluation in dbACC, and finally on outcome processing in dlPFC. In both downstream regions, neural responses to potential and actual rewards decreased as the reference point increased, consistent with predictions from Prospect Theory (1, 33) (Fig. 6A). Together, these results provided direct neural evidence for reference point representation at both single-neuron- and population-level and highlights its broad influence across the decision-making circuit.

**Figure 6:**
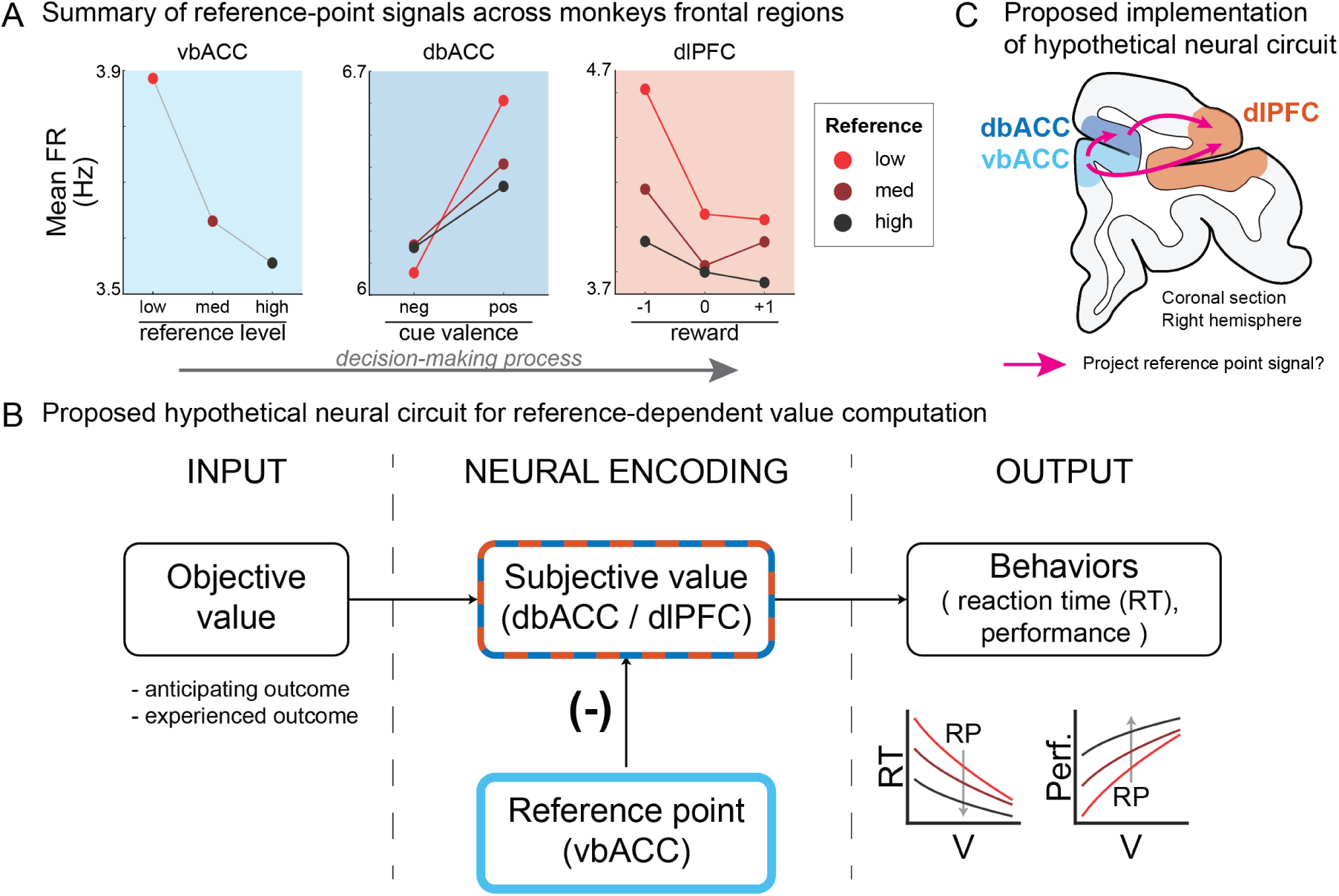
Summary of population-average level reference point signal in monkeys’ frontal areas. A. As the decision-making trial progressed: the vbACC initially tracked the reference point, followed by the dbACC encoding a reference-dependent cue valence signal, and finally, the dlPFC representing the reference-dependent outcome signal. B. A proposed neural circuit for reference-dependent value computation based on the population-level neural activities: the vbACC provides the reference point signal, which modulates decision-variable encoding in the downstream areas, including dbACC and dlPFC. C. Given their anatomical connectivity and reference-related encoding, the vbACC, dbACC, and dlPFC may form an integrated circuit for value computation. Arrows indicate the anatomical directionality of connections.

### A proposed hypothetical neural circuitry for subjective value computation

Based on the distinct population-level encoding patterns observed across monkey frontal regions, we propose a hypothetical neural circuit for computing reference-dependent subjective value. Specifically, we hypothesize that the vbACC serves as a primary source for the reference point signal, providing a global benchmark that modulates value-related activity in downstream regions, including the dbACC and dlPFC (Fig. 6B). The vbACC itself may contain the circuitry required for computing the reference point from environmental and contextual features.

This proposed circuit is consistent with the known anatomical connectivity between these regions. Prior studies have shown that the vbACC projects primarily to both the dbACC and dlPFC (34), and that both vbACC and dbACC project to the dlPFC (35, 36). While our data do not demonstrate how reference point signals are integrated into subjective value, the event-based temporal sequence suggests a potential ventral-to-dorsal progressing direction through this network (Fig. 6C). We note that this circuit is based purely on the anatomical connection, clarifying the exact mechanism of this computational transformation remains a key direction for future research.

### Previous neural evidence of reference point and expectation in the ACC

In a series of theoretical publications, Koszegi and Rabin specifically argued that the reference point should be viewed as “rational expectation” of the environment which complements Prospect Theory by offering a model of how reference points are constructed from reward history (37, 38). In behavioral ecology, the marginal value theorem (MVT) posits that foraging animals should leave a patch when the marginal intake rate falls below the expected average rate of reward in the environment (39). This decision requires animals to track an internal estimate of environmental reward availability, a quantity conceptually and mathematically equivalent to the reference point in Prospect Theory (40). Thus, for many researchers, these instances are mathematically and mechanistically equivalent (41).

Both human and animal studies have identified neural populations that encode these global expected reward rates. In foraging tasks, the ACC encodes a decision variable that signals when to leave the patch in monkeys (15), rodents (17), and humans (16, 18), aligning with our findings on the neuronal origin of the reference point. Consistent with our findings, Hayden and colleagues found that the monkey ACC (including data from regions we identify separately as dbACC and vbACC) encodes a signal correlated with this foraging reference point. This suggests that vbACC activity may reflect a generalized reference point signal across task domains.

More broadly, the ACC has long been implicated in expectation-related processing. It encodes expected outcomes (42–47) and is modulated by both reward history and expectations (11, 48–50). These findings further support the ACC as a neural substrate for reference point representation in non-social decision making. Variability in findings regarding the reference-point (expectation) itself versus reference-dependent encoding may stem from a lack of differentiation between the dorsal and ventral banks of the ACC (above and below the sulcus), which have distinct anatomical structures and functions (51).

### ACC in other decision-making contexts

Beyond non-social decisions, the ACC is also engaged in social and interpersonal decision-making (52, 53), including computing value for both self and others (54, 55). In a closely related study, Chang and colleagues recorded neuronal activity along the dorsal-ventral axis of ACC and found a functional dissociation: encoding of value signals related to others were primarily localized in the ACC gyrus, while the sulcal regions (including regions we identify separately as dbACC and vbACC) contained a mixed representation with neurons encoding either value-to-self, value-to-others or both. This suggests a broader organizational role for the ACC in integrating reference-related value signals across different social contexts.

In social decision-making, however, reference points may extend beyond the internal expectations derived from past outcomes, also including information from external comparisons dependent on social settings. For example, evaluating one’s own salary relative to a colleague’s implicitly involves a social reference point. We speculate that non-social reference points represented in the vbACC may be complemented by social counterparts in the ACC gyrus. Further work is needed and understanding the neural basis of reference point could help extend the current theories of subjective value into social domains.

### Stimulation in vbACC, but not dbACC, changes choice behavior

Few studies have explicitly distinguished between the dorsal and ventral banks of the ACC (14, 20, 54, 56). In one such study, Amemori and Graybiel showed that both regions encode subjective values in monkeys, but the vbACC did so more uniformly and with a stronger negative bias, consistent with our observations. Importantly, microstimulation in the vbACC, but not the dbACC, “increased negative decision-making”, as shown by reduced tolerance toward negative stimuli during an approach-avoidance task. In the context of our study, we hypothesize that stimulating the vbACC is analogous to lowering the reference point, leading to change in subjective value and behaviors in a predictable manner. However, the complexity of their task, which required weighing both positive and negative information for each option, makes it difficult to determine whether this behavioral change was specifically due to a shift in the reference point. Nonetheless, the distinct effects of vbACC versus dbACC stimulation raise specific questions for future research about the unique role of the vbACC in decision-making, specifically encoding the reference point.

### Potential neural mechanisms of reference-dependence

Neural adaptation, change in neural activity in response to changing environmental statistics, is a phenomenon well-documented in sensory systems (57–59). Building on these observations, similar mechanisms have been proposed for value-encoding systems (60, 61). Evidence of range adaptation has been observed across multiple valuation networks, including the dopaminergic system (9, 10, 62); OFC in primates (12, 13); the related area, vmPFC, in human (27, 63); and the ACC in both species (14, 27).

Interestingly, while both OFC and ACC exhibited range adaptation (a context-dependence outcome signal), only the ACC showed a population-level reference point signal in our task. This may reflect several factors, including limited statistical power to detect small effects in the OFC, or the possibility that OFC favors a more mixed representation of value that has been shown to be more flexible for signal encoding (64, 65).

Another possibility is that reference-dependence arises from two distinct neural mechanisms: distributed (intracortical) versus centralized (intercortical) representations of the reference point. In the distributed model, individual brain regions adjust their baselines or tuning curves according to the distribution of values, consistent with adaptation observed in OFC, but without there ever being an explicit representation of the reference point itself. In the centralized model, specialized regions, such as the vbACC, compute the reference point and transmit this signal to downstream value-encoding areas. These regions then integrate both the reference point and values during decision-making (Figs. 1B and 6B). Our findings appear more consistent with the centralized model, while prior work in dopaminergic circuits and the OFC appears to support the distributed form in those areas. Future studies are needed to distinguish these mechanisms more clearly and identify which brain regions operate under each.

### ACC activity as a potential clinical marker of anhedonia and depression

Lastly, our findings offer a possible foundation for future clinical research. Anhedonia, a core feature of major depressive disorder (MDD), reflects a diminished ability to experience positive reinforcement (66, 67). Reference-dependence theories suggest that this may result from a pathologically elevated reference point, which diminishes the perceived value of rewards. Recent studies have proposed that dysfunction in reference point mechanisms may contribute to the emergence of anhedonia in MDD (68, 69).

By localizing the reference point signal to the ACC, our results provide some empirical support for this theoretical hypothesis. Over the past 25 years, Mayberg and colleagues have identified the human subgenual ACC as one of the most consistent neural markers of MDD. Follow-up studies have provided causal evidence, showing that modulating ACC activity can alleviate symptoms in treatment-resistant MDD. Remarkably, recordings from clinical electrodes implanted in the ACC, similar to those presented here, have been shown to correlate with depression severity (70–72).

These findings highlight the ACC’s central role in both value-based decision-making and mental disorders, bridging behavioral economic theory with clinical neuroscience. Understanding how reference point mechanisms in the ACC contribute to mood and motivation may improve our models of MDD and guide the development of more targeted, mechanism-based treatments.

## Conclusion

To summarize, the decisional reference point is a central concept in choice theories, yet its neural instantiation has remained unclear. Our study provides the direct evidence of this signal at the population-level and localizes it to the vbACC. This encoding pattern, along with known anatomical connections to other valuation regions, supports a model of a dedicated neural circuit underlying reference-dependent choice behavior. These findings open new directions for animal studies to test causality through stimulation or pharmacological manipulation, and they offer a clear target (the vbACC) for translational research using fMRI in humans.

## ACKNOWLEDGEMENTS

We thank all colleagues in the Glimcher lab for their support and helpful comments. This work was supported by the NIMH T32 Training program to D.N., National Institutes of Health - Office of Extramural Research Grants R01DA19028, P01NS040813 and R01-MH132640 to J.D.W., R01MH121480 and a Hilda and Preston Davis Foundation grant to E.L.R., and NIH U19 NS107616 to K.L..

## AUTHOR CONTRIBUTIONS

Conceptualization - ELR, JDW, and PWG

Data curation - ELR and JDW

Formal analysis - DN, KL, and PWG

Funding acquisition - JDW and PWG

Investigation - ELR and JDW

Methodology - ELR and JDW

Project administration - ELR, JDW, KL, and PWG

Resources - ELR and JDW

Software - DN and ELR

Supervision - ELR, KL, and PWG

Validation - DN, ELR, JDW, KL, and PWG

Visualization - DN

Writing – original draft - DN, KL, and PWG

Writing – review & editing – DN, ELR, JDW, KL, and PWG

## DATA AND CODE AVAILABILITY

Preprocessed data and custom code is available at OSF directory.

## METHODS

### Subjects and Behavioral Task

The subjects were two male rhesus macaques (Macaca mulatta), named M and C, aged 6 and 10 years old, and weighing 11 and 14.5 kg at the time of recording, respectively. The monkeys were trained to perform an instructed response task, where they moved a bidirectional joystick left or right in response to visual cues displayed at the center of a computer screen. The task involved four different cues, differing by correct response direction (left/right) and valence (positive/negative) in a 2×2 design. Two cues (one left and one right) could lead to an increased earned reward if the response was correct, and no change if incorrect (positive valence). The other two cues (one left and one right) resulted in a reduced earned reward if the response was incorrect, and no change if correct (negative valence) (Fig. 2B). Each cue represented a unique combination of valence and response instruction. The same four visual cues were used for each monkey across all sections. For every trial, a cue was chosen purely randomly and independent of other trials. Changes in the earned rewards were reflected by adjustments in the length of the accumulated-reward bar displayed at the bottom of the screen. The magnitude of the increases and decreases was equal and fixed across trials. To ensure that sufficient trials resulted in negative feedback (equivalent to error response), 15% of correct responses ended in the worse outcome (no change for positive cue trials; decreased earned reward for negative cue trials).

The subjects completed six trials within each block before the accumulated reward earned over the last six trials was delivered (Fig. 2C). This number of trials in a block was calibrated to maintain the subjects’ motivation throughout the block while maximizing variation in the accumulated-reward bar length. Rewards were fruit juice, and the amount received was directly proportional to the length of the accumulated-reward bar at the end of the trial block. A trial was considered complete when the subject made a joystick response to the visual stimulus, regardless of the outcome. Incomplete trials, where the monkey failed to make a joystick response or failed to successfully hold initial fixation, did not count toward block completion. After juice delivery, the accumulated-reward bar length was reset to a value of 2 at the start of the next block.

Behavioral contingencies were implemented using MonkeyLogic software (73), and eye movements were tracked with an infrared video-based system (iSCAN). Each subject was surgically implanted with two recording chambers and a titanium head post to stabilize the head position during recording. The precise implantation sites were determined based on stereotaxic coordinates of target areas identified on 1.5T MRI scans of each subject’s brain. All procedures were carried out following the guidelines of the National Institutes of Health Guide for the Care and Use of Laboratory Animals and the recommendations of the University of California at Berkeley Animal Care and Use Committee.

Each recording session consisted of hundreds of consecutive trials (average 646.90 ± 12.43). At the start of each trial, only the accumulated-reward bar that reflected the accumulated reward was visible, followed by a fixation point at the center of the screen. Subjects were required to hold fixation within a 1.3° radius of the fixation point for 650 ms to initiate the trial. After fixation, one of four visual cues, which were images of natural scenes approximately 2°×3° in size, was presented. Subjects responded by moving a joystick fixed to the front of their chair either left or right. To discourage random responses, any response made within 150 ms of stimulus onset triggered a 5-second timeout. After the response, feedback was provided through changes in the length of the accumulated-reward bar (Fig. 2A).

In this setup, the accumulated-reward bar remained visible throughout each trial and its length carried over from one trial to the next within a block, so that the accumulated-reward bar consistently indicated the total amount of reward the subjects had accumulated and served as the reference point in classic economic decision-making. This design ensured that subjects were continuously aware of their current earned reward. Additionally, by fixing the amounts of reward increases and decreases, we controlled the magnitude of gains and losses, enabling us to examine the effect of reference levels without confounding subjective perceptions of value.

### Neural recording and general processing

Neural recordings were acquired using a Plexon MAP system, with 4 to 20 tungsten microelectrodes (FHC Instruments), independently positioned in the targeted brain regions during each session (see Fig. 2E). The targeted brain regions were broadly labeled as follows: vbACC (areas 24a & 24b), dbACC (areas 24b & 24c), mOFC (area 14), OFC (area 13), dlPFC (areas 9, 46, & 9/46), and vlPFC (area 47/12). Precisely, our dbACC (vbACC) target was the dorsal (ventral) bank of the cingulate sulcus within the rostral (pre and perigenual) ACC. Areal boundaries were defined by sulci visible on MRIs, as shown in Fig 2E. During recording, electrodes were placed in cortical layers by acoustic mapping of gray-white matter boundaries of each electrode path. After recording, final placement was determined on MR images for each subject using stereotaxic coordinates and depth of each electrode. If a neuron could not be confidently assigned to one of the anatomical locations, it was excluded from further analysis.

All well-isolated units were recorded, with no pre-screening. The recorded waveforms were digitized, and single units were identified using Offline Sorter (Plexon). After spike sorting, we excluded units with an average firing rate below 1 Hz across the entire session, as these neurons did not show sufficient activity for reliable statistical analysis. The final number of neurons used for analysis in each brain region was: dbACC (n = 154), vbACC (n = 156), mOFC (n = 126), OFC (n = 170), dlPFC (n = 159), and vlPFC (n = 212).

### Behavioral analysis

Behavioral performance, including trial-by-trial percent correct and reaction time, across all sessions and monkeys were aggregated and analyzed using a mixed-effects logistic regression and a mixed-effects regression model, respectively. We started with models including only the reference point, cue valence and their interaction:

For reaction time:

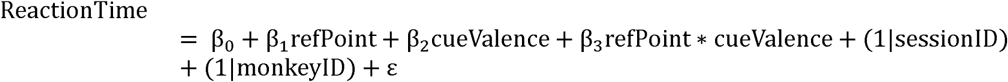

For percent correct (PC):

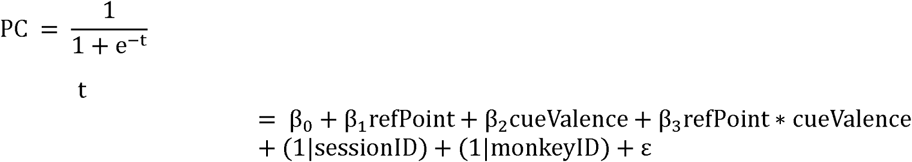

where *refPoint* is the reference point; *cueValence* is the cue valence; and their interaction term; *sessionID* is recorded session indicator; *monkeyID* is subject indicator.

To control for potential confounds, we expanded above models to include more regressors that are potentially covaried with the reference point:

For reaction time:

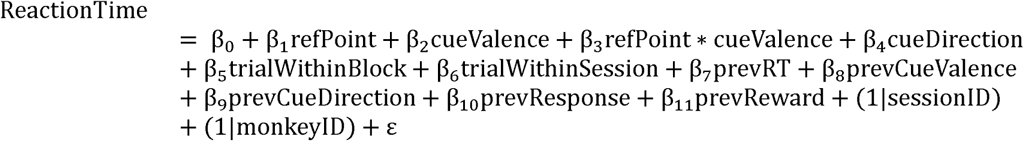

For percent correct (PC):

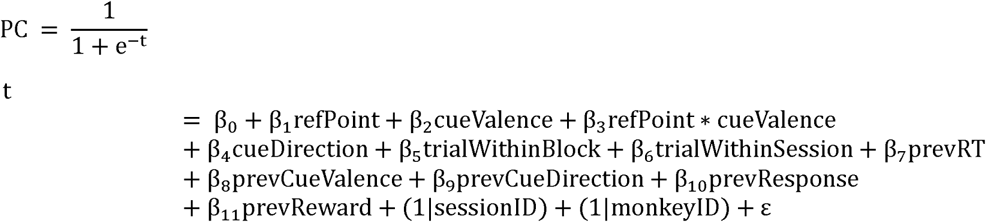

where cueDirection is the response-instruction of the cue; trialWithinBlock is the current trial’s position within the block (1–6); trialWithinSession is the current trial’s position in the session to control of drifting performance over time; prevRT, prevCueValence, prevCueDirection, prevResponse and prevReward are reaction-time, cue-valence, response-instruction, response (correct/incorrect) and outcome (−1/0/+1) in the previous trial.

### Single-unit regression

#### Primary analyses in three main epochs

For each neuron, we calculated its firing rate during three event epochs: *trial start*, *cue onset*, and *feedback*. For each epoch, spikes were aligned to the event onset for all trials, we then averaged spike counts in 150 ms time bins stepped at 25ms, with bin centers ranging from 200ms preceding to 650 ms following event onset. We analyzed neural activities within a 500ms time window beginning 100ms after event onset (101-600ms post-event; Fig. 3A). This window was chosen a priori based on the reaction time distribution in the cue onset epoch, which was selected to approximately capture the time preceding median responses of both subjects. The same analysis window was used for all epochs. Only completed trials were included in the analysis.

Within this window, we averaged spike count for mean firing rate (mean FR), a standard calculation in previous studies (19, 20). We used this activity to fit multivariable regression models for each neuron for each epoch. Models used are as follows:

- Trial start epoch:

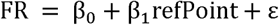

where ε is the error term, *refPoint* corresponds to length of the accumulated-reward bar at the beginning of each trial. We reported population analysis for β_1_ in the main text.
- Cue onset epoch:

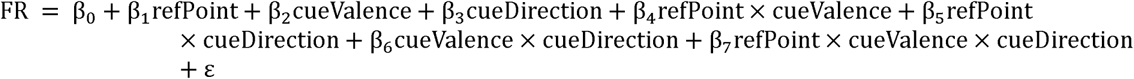

where ε is the error term, *refPoint* corresponds to length of the accumulated-reward bar at the beginning of each trial; *cueValence* and *cueDirection* corresponds to valence (positive/ negative) and response instruction (left/ right) of the cue of each trial. We reported population analysis for β_4_ in the main text.
- Feedback epoch:

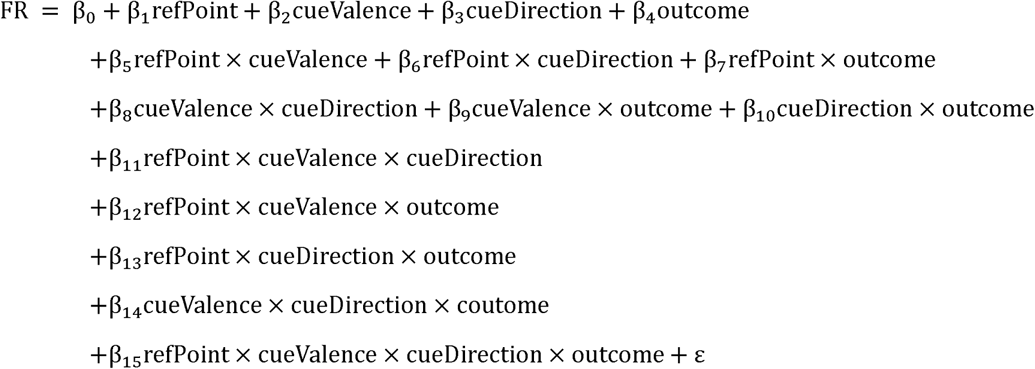

where ε is the error term, *refPoint* corresponds to length of the accumulated-reward bar at the beginning of each trial; *cueValence* and *cueDirection* corresponds to valence (positive/ negative) and response instruction (left/ right) of the cue of each trial; and *outcome* refers to earned-reward/ change in accumulated-reward-bar (+1/0/−1). We reported population analysis for β_7_ in the main text.

The model coefficient is influenced by the firing rate profile of each neuron, which can vary with firing rate range and response timing across populations. In addition to mean FR, we also calculated other types of firing rates, including: (1) standardized mean FR, where the mean FR of each neuron was normalized based on the variance of these values across all trials within a session before fitting the model; (2) peak FR, which is the peak firing rate in a 150 ms window within a 600ms time window beginning right after event onset (1-600ms post-event); and (3) standardized peak FR, where peak FR was normalized based on the variance of these values across all trials within a session before fitting the model (see Fig. S2A). Peak FR was included to account for neurons with different response delays, where the mean FR in the analysis window could be higher or lower due to earlier or later responses. Standardizing these FR values prevents impact of outliers when analyzing population-level statistics.

### Latent dimension analysis

#### Principal Component Analysis (PCA)

Neural activities were aggregated to into pseudo-populations separately for each brain region. As described above, spike counts were aligned to the onset of three epochs: trial-start, cue-onset and feedback. For each epoch, we generated a *t1 × c × n* matrix:

- *t1* is the number of time points (101-600ms post-event)
- *c* is the number of conditions

- trial-start: 6 conditions (<=1, 2-5, or >= 6 bar-length)
- cue-onset: 6 conditions (3 reference-levels *×* 2 cue-valence)
- feedback: 9 conditions (3 reference-levels *×* 3 outcome levels)
- *n* is the number of neurons per brain region

For each region, computed the condition-averaged activity across neuron, resulting in an *n × c* matrix. PCA was applied on this matrix using MATLAB’s *pca* function to extract latent dimensions that captured the dominant variance in each area. These dimensions represent the neural latent space that explains reference-related information within each group of neurons. Coefficients of each condition in this space are plotted in the left columns of supplemental figures (Figs. S1, S6, S8)

To visualize the temporal dynamics, we generated a *t2 × c × n* matrix, where *t2* is the number of time points ranging from 200ms pre- to 1000ms post-event. Neural activity for each condition was projected onto the first, second and third PCs, and temporal trajectories were plotted to show condition-specific dynamics across time in each brain region (Figs. S1, S6, S8, middle columns).

#### Demixed Principal Component Analysis (dPCA)

To dissociate task-related factors from general temporal structure, we also performed dPCA. Using the same *t2 × c × n* matrix described above, dPCA was applied following the same approach described earlier (26). dPCA extracted “interpretable” dimensions associated with conditions and time specifically. Temporal trajectories in the first three demixed PCs were shown in the right columns of supplemental figures (Figs. S1, S6, S8), demonstrating the reference-related signals in the demixed latent space.

### Population-level analysis

To assess a brain region’s specific response to a model regressor, we examined the β*s* (coefficients) and the corresponding *p-value*s (Fig. 3A) of all neurons recorded in that area, encompassing six different areas recorded across all recording sessions (Fig. 2E).

#### Primary analysis

To assess whether a brain area exhibited homogeneous, population-level, responses to a specific decision variable, we performed two complementary analyses:

1. Mean regression coefficient analysis. We evaluated the mean ± SEM of the coefficients (β values) for all neurons in each area. A two-tailed one-sample *t*-test against zero was used to determine whether the average β significantly differed from zero. A significant result indicated a consistent, population-level modulation by the decision variable.
2. Proportion of tuned neurons analysis. We calculated the percentage of the population (PoP) with significant coefficients (p < 0.05) for each area for the variable of interest. These neurons were then categorized by the sign of their β (positive or negative). We calculated the proportion of positive and negative neurons as:

- Positive PoP: 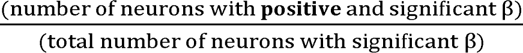
- Negative PoP: 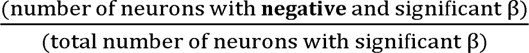

A two-tailed χ^2^ test was used to determine whether there was a significant bias in the percentage of positive versus negative populations.

To account for multiple comparisons across the six brain regions, we applied both uncorrected (p < 0.05) and Bonferroni-corrected significance thresholds for six brain regions (p < 0.0083).

#### Controlling for differences across recorded sessions and subjects

To ensure that our findings were not biased by differences between subjects, we applied mixed-effects models using sessionID and monkeyID as random effects:

1. Mean regression coefficient analysis. For each brain region, we used a linear mixed-effects model to test whether the mean regression coefficient (β) significantly deviated from zero while accounting for subject variability.

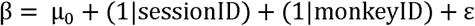
2. Proportion of tuned neurons analysis. For each brain region, we identified neurons with significant tuning (p < 0.05) and categorized them by the sign of their β coefficient (positive or negative). We then applied a logistic mixed-effects model to test whether the proportion of positively tuned neurons deviated from chance, again accounting for subject as a random effect.

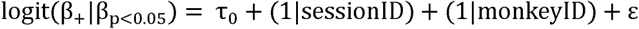

#### Controlling for potential confounds

To account for confounding factors that may covary with the reference point, we applied a hierarchical modeling approach. For **each epoch and each neuron**, we first regressed out potential confounds, including:

- *trialWithinBlock* – trial number within a block (1–6)
- *trialWithinSession* – trial number within the session (control for performance drift)
- *prevCueValence* – valence of previous trial’s cue
- *prevCueDirection* – response instruction on previous trial
- *prevResponse* – correctness of previous trial’s response
- *prevReward* – outcome of the previous trial (−1/0/+1)

We then computed the residuals from these models, defined as the difference between real and predicted firing rate, and used them in the main regression models to isolate the effect of the reference point (see “Primary Analysis” section). Specifically:

- Trial start epoch:

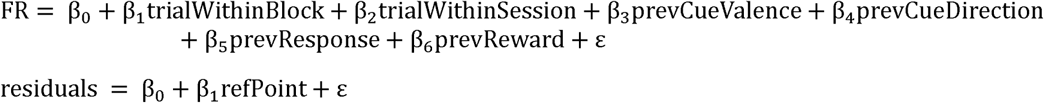
- Cue onset epoch:

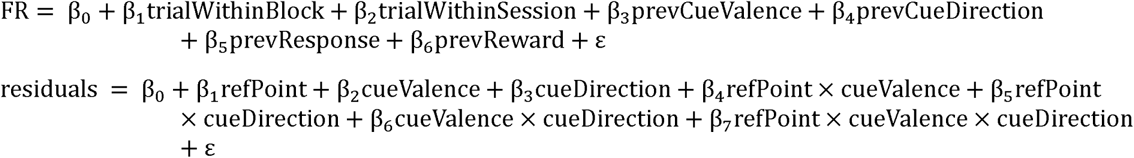
- Feedback epoch:

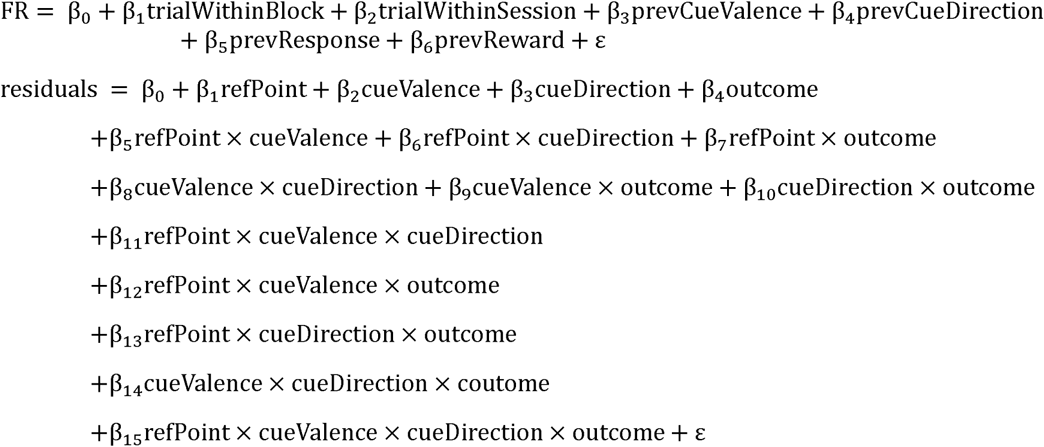

We reported the relevant coefficients of interest in the supplementary figures: β_1_ in the trial start epoch (Fig. S4), β_4_ in the cue onset epoch (Fig. S7), and β_7_ in the feedback epoch (Fig. S9).

Finally, we combined this hierarchical control with the mixed-effects models described in the “Controlling for Subject Differences” section.

### Population-level Neural Dynamics

For each neuron in each event epoch, spikes were aligned to the event onset for all trials, we then averaged spike counts in 150 ms time bins stepped at 25ms, with bin centers ranging from 1000ms preceding cue presentation to 1000 ms following event onset. We then grouped trials into 3 reference-level categories (Fig. 2D) and calculated the mean FR in each one.

For all neurons in each brain area, we continued averaging FR across all neurons in each reference-level category, resulting in mean ± SEM for each time bin.

### Further analyses for vbACC

#### Testing the lasting effect of reference point in vbACC throughout trial

First, using the population neural dynamics described above, we calculated the average FR within a 500ms time window beginning 100ms after event onset (for fixation, it is the fixation start) for different reference levels. We then performed repeated-measurement ANOVA for each epoch to test the difference of neural activity due to reference levels (Fig. S5B).

Second, for each neuron in the vbACC, we calculated its firing rate during five event epochs: trial start, fixation, cue onset, response, and feedback. For each epoch, analyses were done as described in “Single-unit regression” section but using only univariate regression with *refPoint* as the sole regressor. Figure S5C showed the result of population analysis for these univariate regressions.

### General statistics

P-values less than or equal to 0.05 were considered significant (*p < 0.05, **p < 0.01, ***p < 0.001, n.s. p > 0.05). Bonferroni correction for six brain regions was applied to control for multiple comparisons (‡p ≤ 0.0083). All figures show mean ± SEM, except otherwise noted.

