## Supplemental information for "Neural representation of the decisional reference point in monkeys"

**Supplemental Information File**

### This file includes:

### Supplemental Tables S1-S3:

**For Tables S1-3:**

Coefficients – Coefficients of regression model corresponding to the variable

- Mean / SE – mean and standard error of coefficient values for all neurons
- P-value: p-value of one-sample t-test for all neurons in each brain region

PoP – The percentage of population with significant coefficient values for all regressors.

- %-pos: percentage of population with significantly positive
- %-neg: percentage of population with significantly negative
- P-value: p-value of $\chi^2$ test for positive versus negative for each brain region

Shaded rows are the main effects reported in the main text.

**Table S1**: Statistics of population encoding of the reference point during trial start epoch (Supplemental for Fig 3)

| | | dbACC | dlPFC | vlPFC | mOFC | OFC | vbACC |
|---|---|---|---|---|---|---|---|
| Reference point | Coefficients<br>Mean<br>(SE)<br>p-value | 0.0456<br>(0.0528)<br>0.3890 | 0.0152<br>(0.0387)<br>0.6955 | 0.0054<br>(0.0410)<br>0.8951 | -0.0292<br>(0.0309)<br>0.3466 | -0.0446<br>(0.0267)<br>0.0968 | -0.1050<br>(0.0374)<br>0.0056 |
| | PoP<br>%-pos<br>%-neg<br>p-value | 25.32<br>29.87<br>0.2829 | 16.35<br>22.01<br>0.1032 | 18.87<br>27.83<br>0.0069 | 9.52<br>19.84<br>0.0017 | 14.12<br>24.71<br>0.0017 | 10.26<br>27.56<br>$6.66\times10^{-7}$ |

**Table S2**: Statistics of population encoding during cue onset epoch (Supplemental for Fig. 4)

| | | dbACC | dlPFC | vlPFC | mOFC | OFC | vbACC |
|---|---|---|---|---|---|---|---|
| Reference point (RP) | Coefficients<br>Mean<br>(SE)<br>p-value | 0.0688<br>0.0681<br>0.3141 | 0.0098<br>0.0419<br>0.8151 | 0.0082<br>0.0498<br>0.8697 | -0.0114<br>0.0295<br>0.7000 | -0.0146<br>0.0392<br>0.7110 | -0.0280<br>0.0384<br>0.4666 |
| | PoP<br>%-pos<br>%-neg<br>p-value | 18.18<br>16.88<br>0.7003 | 13.83<br>15.72<br>0.5360 | 10.84<br>10.37<br>0.8330 | 5.55<br>7.14<br>0.4795 | 10.00<br>11.17<br>0.6374 | 7.69<br>8.33<br>0.7773 |
| Cue valence (Val) | Coefficients<br>Mean<br>(SE)<br>p-value | 0.8947<br>0.3459<br>0.0106 | -0.2765<br>0.2159<br>0.2022 | 0.5354<br>0.3544<br>0.1323 | 0.0282<br>0.1607<br>0.8612 | -0.0344<br>0.1726<br>0.8422 | -0.1535<br>0.1096<br>0.1634 |
| | PoP<br>%-pos<br>%-neg<br>p-value | 21.43<br>14.29<br>0.0359 | 5.03<br>15.72<br>$2.85 \times 10^{-5}$ | 16.51<br>13.68<br>0.2888 | 7.14<br>9.52<br>0.3545 | 11.18<br>12.35<br>0.6547 | 5.13<br>6.41<br>0.5050 |
| Cue direction instruction (Dir) | Coefficients<br>Mean<br>(SE)<br>p-value | 0.1701<br>0.1947<br>0.3838 | 0.0495<br>0.1805<br>0.7845 | -0.0845<br>0.1095<br>0.4412 | -0.0987<br>0.1064<br>0.3551 | 0.1789<br>0.0886<br>0.0451 | 0.0508<br>0.1058<br>0.6316 |
| | PoP<br>%-pos<br>%-neg<br>p-value | 9.74<br>6.49<br>0.1573 | 6.91<br>6.91<br>1.0000 | 8.01<br>5.66<br>0.1892 | 4.76<br>3.17<br>0.3711 | 4.11<br>2.94<br>0.4142 | 3.20<br>2.56<br>0.6374 |
| RP × Val | Coefficients<br>Mean<br>(SE)<br>p-value | -0.1970<br>0.0593<br>0.0011 | 0.0338<br>0.0452<br>0.4564 | -0.0606<br>0.0538<br>0.2609 | -0.0290<br>0.0354<br>0.4143 | -0.0071<br>0.0357<br>0.8416 | 0.0067<br>0.023<br>0.8191 |
| | PoP<br>%-pos<br>%-neg<br>p-value | 6.49<br>14.94<br>0.0014 | 10.06<br>6.29<br>0.0961 | 6.13<br>8.96<br>0.1336 | 7.14<br>3.97<br>0.1306 | 7.65<br>5.29<br>0.2278 | 4.49<br>1.28<br>0.0184 |
| RP × Dir | Coefficients<br>Mean<br>(SE)<br>p-value | -0.0521<br>0.0487<br>0.2866 | -0.0001<br>0.0374<br>0.9969 | 0.0431<br>0.0343<br>0.2104 | 0.0325<br>0.0315<br>0.3033 | -0.0554<br>0.0293<br>0.0599 | 0.0096<br>0.0310<br>0.7566 |
| | PoP<br>%-pos<br>%-neg<br>p-value | 5.84<br>8.44<br>0.2278 | 5.03<br>4.40<br>0.7150 | 6.60<br>2.83<br>0.0114 | 1.58<br>2.38<br>0.5271 | 3.52<br>4.70<br>0.4497 | 1.92<br>1.28<br>0.5271 |
| Val × Dir | Coefficients<br>Mean<br>(SE)<br>p-value | -0.2405<br>0.2337<br>0.3051 | 0.1664<br>0.1793<br>0.3547 | -0.1483<br>0.1790<br>0.4084 | 0.1153<br>0.1481<br>0.4377 | -0.0778<br>0.1527<br>0.6111 | -0.0343<br>0.1473<br>0.8162 |
| | PoP<br>%-pos<br>%-neg<br>p-value | 5.84<br>13.63<br>0.0019 | 6.28<br>4.40<br>0.3035 | 7.07<br>10.37<br>0.1036 | 2.38<br>2.38<br>1.0000 | 5.29<br>4.11<br>0.4795 | 1.28<br>1.92<br>0.5271 |
| RP × Val × Dir | Coefficients<br>Mean<br>(SE)<br>p-value | 0.1058<br>0.0596<br>0.0778 | -0.0153<br>0.0489<br>0.7553 | 0.0083<br>0.0448<br>0.8523 | -0.0110<br>0.0423<br>0.7945 | 0.0289<br>0.0398<br>0.4685 | 0.0165<br>0.0430<br>0.7021 |
| | PoP<br>%-pos<br>%-neg<br>p-value | 9.09<br>1.94<br>0.0002 | 3.77<br>3.77<br>1.0000 | 7.07<br>3.30<br>0.0159 | 1.58<br>1.58<br>1.0000 | 2.94<br>3.53<br>0.6698 | 3.20<br>1.28<br>0.1088 |

**Table S3**: Statistics of population encoding during feedback epoch (Supplemental for Fig. 5).

| [redacted] | | dbACC | dlPFC | vlPFC | mOFC | OFC | vbACC |
|---|---|---|---|---|---|---|---|
| Reference point (RP) | Coefficients<br>Mean<br>(SE)<br>p-value | -0.0589<br>0.0688<br>0.3932 | -0.1122<br>0.0538<br>0.0389 | -0.0324<br>0.0520<br>0.5336 | -0.0175<br>0.0506<br>0.7299 | -0.0401<br>0.0359<br>0.2652 | -0.0986<br>0.0354<br>0.0059 |
| | PoP<br>%-pos<br>%-neg<br>p-value | 18.18<br>20.13<br>0.5807 | 5.03<br>13.83<br>0.0003 | 13.20<br>17.45<br>0.1144 | 7.14<br>10.31<br>0.2278 | 5.29<br>10.59<br>0.0143 | 7.69<br>11.54<br>0.1213 |
| Cue valence (Val) | Coefficients<br>Mean<br>(SE)<br>p-value | 0.5369<br>0.3553<br>0.1328 | -0.0082<br>0.2491<br>0.9739 | 0.5378<br>0.2887<br>0.0639 | 0.0051<br>0.2324<br>0.9826 | -0.2168<br>0.2099<br>0.3031 | 0.2084<br>0.2161<br>0.3363 |
| | PoP<br>%-pos<br>%-neg<br>p-value | 14.93<br>9.09<br>0.0364 | 11.32<br>8.80<br>0.3173 | 11.32<br>9.90<br>0.5271 | 7.93<br>6.34<br>0.5050 | 4.70<br>9.41<br>0.0209 | 7.69<br>3.84<br>0.0455 |
| Cue direction instruction (Dir) | Coefficients<br>Mean<br>(SE)<br>p-value | 0.3235<br>0.1586<br>0.0431 | -0.0708<br>0.1716<br>0.6805 | 0.1908<br>0.1690<br>0.2601 | 0.1052<br>0.1119<br>0.3490 | 0.1107<br>0.1171<br>0.3459 | 0.1316<br>0.1310<br>0.3167 |
| | PoP<br>%-pos<br>%-neg<br>p-value | 6.49<br>3.89<br>0.1573 | 6.28<br>6.91<br>0.7576 | 9.43<br>5.18<br>0.0223 | 5.55<br>3.96<br>0.4142 | 3.52<br>3.52<br>1.0000 | 7.69<br>4.48<br>0.1048 |
| Outcome (Rew) | Coefficients<br>Mean<br>(SE)<br>p-value | 0.2012<br>0.3124<br>0.5204 | -0.9745<br>0.2961<br>0.0012 | 0.0088<br>0.2325<br>0.9697 | -0.2222<br>0.2221<br>0.3192 | -0.2781<br>0.1892<br>0.1436 | -0.2943<br>0.1785<br>0.1013 |
| | PoP<br>%-pos<br>%-neg<br>p-value | 9.09<br>9.74<br>0.7928 | 5.03<br>20.12<br>$8.02\times10^{-8}$ | 8.01<br>6.60<br>0.4461 | 2.38<br>10.31<br>$4.07\times10^{-4}$ | 2.94<br>5.29<br>0.1306 | 2.56<br>5.76<br>0.0499 |
| RP × Val | Coefficients<br>Mean<br>(SE)<br>p-value | -0.0345<br>0.0698<br>0.6216 | 0.0488<br>0.0658<br>0.4596 | -0.0544<br>0.0628<br>0.3873 | -0.0235<br>0.0629<br>0.7094 | 0.0691<br>0.0552<br>0.2128 | -0.0468<br>0.0656<br>0.4773 |
| | PoP<br>%-pos<br>%-neg<br>p-value | 7.79<br>10.38<br>0.2850 | 9.43<br>6.91<br>0.2673 | 7.54<br>9.43<br>0.3458 | 6.34<br>7.14<br>0.7316 | 5.88<br>2.94<br>0.0679 | 3.20<br>4.48<br>0.4142 |
| RP × Dir | Coefficients<br>Mean<br>(SE)<br>p-value | -0.0287<br>0.0457<br>0.5310 | -0.0036<br>0.0488<br>0.9420 | -0.0897<br>0.0447<br>0.0459 | -0.0260<br>0.0373<br>0.4870 | -0.0354<br>0.0363<br>0.3302 | -0.0093<br>0.0347<br>0.7879 |
| | PoP<br>%-pos<br>%-neg<br>p-value | 6.49<br>5.19<br>0.5050 | 7.54<br>5.03<br>0.2059 | 2.35<br>8.49<br>0.0001 | 3.96<br>4.76<br>0.6698 | 4.70<br>2.35<br>0.1025 | 4.48<br>4.48<br>1.0000 |
| RP × Rew | Coefficients<br>Mean<br>(SE)<br>p-value | -0.0824<br>0.0938<br>0.3813 | 0.1993<br>0.0831<br>0.0176 | -0.0276<br>0.0735<br>0.7083 | 0.0353<br>0.0663<br>0.5950 | 0.0563<br>0.0658<br>0.3931 | 0.0048<br>0.0547<br>0.9297 |
| | PoP<br>%-pos<br>%-neg<br>p-value | 7.14<br>7.79<br>0.7681 | 12.57<br>4.40<br>0.0004 | 7.07<br>7.07<br>1.0000 | 9.52<br>3.17<br>0.0047 | 4.11<br>5.88<br>0.3035 | 3.20<br>2.56<br>0.6374 |

| | | | | | | | |
|---|---|---|---|---|---|---|---|
| Val × Dir | Coefficients<br>Mean<br>(SE)<br>p-value | -0.2567<br>0.2797<br>0.3602 | -0.3008<br>0.2695<br>0.2660 | -0.0385<br>0.2371<br>0.8713 | -0.4714<br>0.2117<br>0.0278 | 0.0095<br>0.2069<br>0.9636 | -0.4194<br>0.2510<br>0.0967 |
| | PoP<br>%-pos<br>%-neg<br>p-value | 3.89<br>5.19<br>0.4497 | 4.40<br>4.40<br>1.0000 | 3.30<br>3.77<br>0.7150 | 1.58<br>3.96<br>0.1088 | 1.76<br>3.52<br>0.1573 | 0.64<br>4.48<br>0.0027 |
| Val × Rew | Coefficients<br>Mean<br>(SE)<br>p-value | -0.1051<br>0.3392<br>0.7572 | 0.9182<br>0.3823<br>0.0175 | -0.1457<br>0.3192<br>0.6485 | 0.1996<br>0.2843<br>0.4839 | 0.4665<br>0.2168<br>0.0329 | -0.1020<br>0.2537<br>0.6882 |
| | PoP<br>%-pos<br>%-neg<br>p-value | 8.44<br>7.79<br>0.7773 | 11.32<br>3.77<br>0.0005 | 8.49<br>8.49<br>1.0000 | 3.96<br>3.96<br>1.0000 | 4.70<br>1.76<br>0.0330 | 5.12<br>3.84<br>0.4497 |
| Dir × Rew | Coefficients<br>Mean<br>(SE)<br>p-value | 0.5417<br>0.3606<br>0.1351 | 0.2895<br>0.3956<br>0.4654 | -0.0329<br>0.3174<br>0.9176 | 0.6259<br>0.3008<br>0.0395 | 0.2358<br>0.2286<br>0.3038 | 0.1393<br>0.2753<br>0.6137 |
| | PoP<br>%-pos<br>%-neg<br>p-value | 7.79<br>3.89<br>0.0455 | 10.06<br>5.03<br>0.0209 | 4.24<br>6.13<br>0.2278 | 8.73<br>3.17<br>0.0106 | 5.29<br>4.11<br>0.4795 | 5.76<br>3.84<br>0.2733 |
| RP × Val × Dir | Coefficients<br>Mean<br>(SE)<br>p-value | 0.0539<br>0.0801<br>0.5018 | 0.0675<br>0.0864<br>0.4360 | 0.0386<br>0.0717<br>0.5905 | 0.1502<br>0.0662<br>0.0250 | -0.0238<br>0.0640<br>0.7104 | 0.1242<br>0.1043<br>0.2358 |
| | PoP<br>%-pos<br>%-neg<br>p-value | 3.89<br>3.89<br>1.0000 | 3.77<br>3.77<br>1.0000 | 4.71<br>3.30<br>0.3035 | 3.96<br>1.58<br>0.1088 | 1.76<br>2.35<br>0.5930 | 5.76<br>0.64<br>0.0003 |
| RP × Val × Rew | Coefficients<br>Mean<br>(SE)<br>p-value | 0.0105<br>0.0973<br>0.9143 | -0.2099<br>0.1031<br>0.0434 | 0.0090<br>0.0987<br>0.9274 | -0.0155<br>0.0860<br>0.8571 | -0.1279<br>0.0772<br>0.0992 | 0.0023<br>0.0762<br>0.9763 |
| | PoP<br>%-pos<br>%-neg<br>p-value | 7.14<br>4.54<br>0.1824 | 2.51<br>10.69<br>0.0001 | 7.07<br>6.60<br>0.7928 | 2.38<br>5.55<br>0.0736 | 3.52<br>6.47<br>0.0863 | 1.92<br>3.20<br>0.3173 |
| RP × Dir × Rew | Coefficients<br>Mean<br>(SE)<br>p-value | -0.1163<br>0.1128<br>0.3039 | -0.0781<br>0.1122<br>0.4876 | 0.0027<br>0.0970<br>0.9780 | -0.2213<br>0.0960<br>0.0227 | -0.0735<br>0.0742<br>0.3235 | -0.0289<br>0.0905<br>0.7497 |
| | PoP<br>%-pos<br>%-neg<br>p-value | 5.19<br>6.49<br>0.5050 | 3.14<br>6.28<br>0.0679 | 3.77<br>3.77<br>1.0000 | 2.38<br>6.34<br>0.0330 | 1.76<br>2.94<br>0.3173 | 2.56<br>2.56<br>1.0000 |
| Val × Dir × Rew | Coefficients<br>Mean<br>(SE)<br>p-value | -0.6225<br>0.4015<br>0.1231 | 0.0748<br>0.4373<br>0.8644 | 0.0035<br>0.3964<br>0.9929 | -0.1042<br>0.3516<br>0.7675 | -0.2738<br>0.3068<br>0.3733 | 0.0688<br>0.3289<br>0.8346 |
| | PoP<br>%-pos<br>%-neg<br>p-value | 3.89<br>4.54<br>0.6949 | 3.14<br>6.91<br>0.0339 | 4.71<br>5.66<br>0.5465 | 4.76<br>1.58<br>0.0455 | 1.17<br>4.70<br>0.0073 | 2.56<br>2.56<br>1.0000 |
| RP × Val × Dir × Rew | Coefficients<br>Mean<br>(SE)<br>p-value | 0.1222<br>0.1238<br>0.3252 | -0.0247<br>0.1278<br>0.8470 | 0.0175<br>0.1210<br>0.8848 | 0.0280<br>0.1170<br>0.8112 | 0.1350<br>0.0972<br>0.1667 | -0.0144<br>0.1189<br>0.9041 |
| | PoP | 3.89 | 6.28 | 3.77 | 1.58 | 4.11 | 1.92 |

| | | | | | | | |
|---|---|---|---|---|---|---|---|
| | %-pos<br>%-neg<br>p-value | 3.24<br>0.6698 | 2.51<br>0.0233 | 2.83<br>0.4497 | 3.17<br>0.2482 | 1.17<br>0.0184 | 2.56<br>0.5930 |

### Supplemental Figures S1-S9

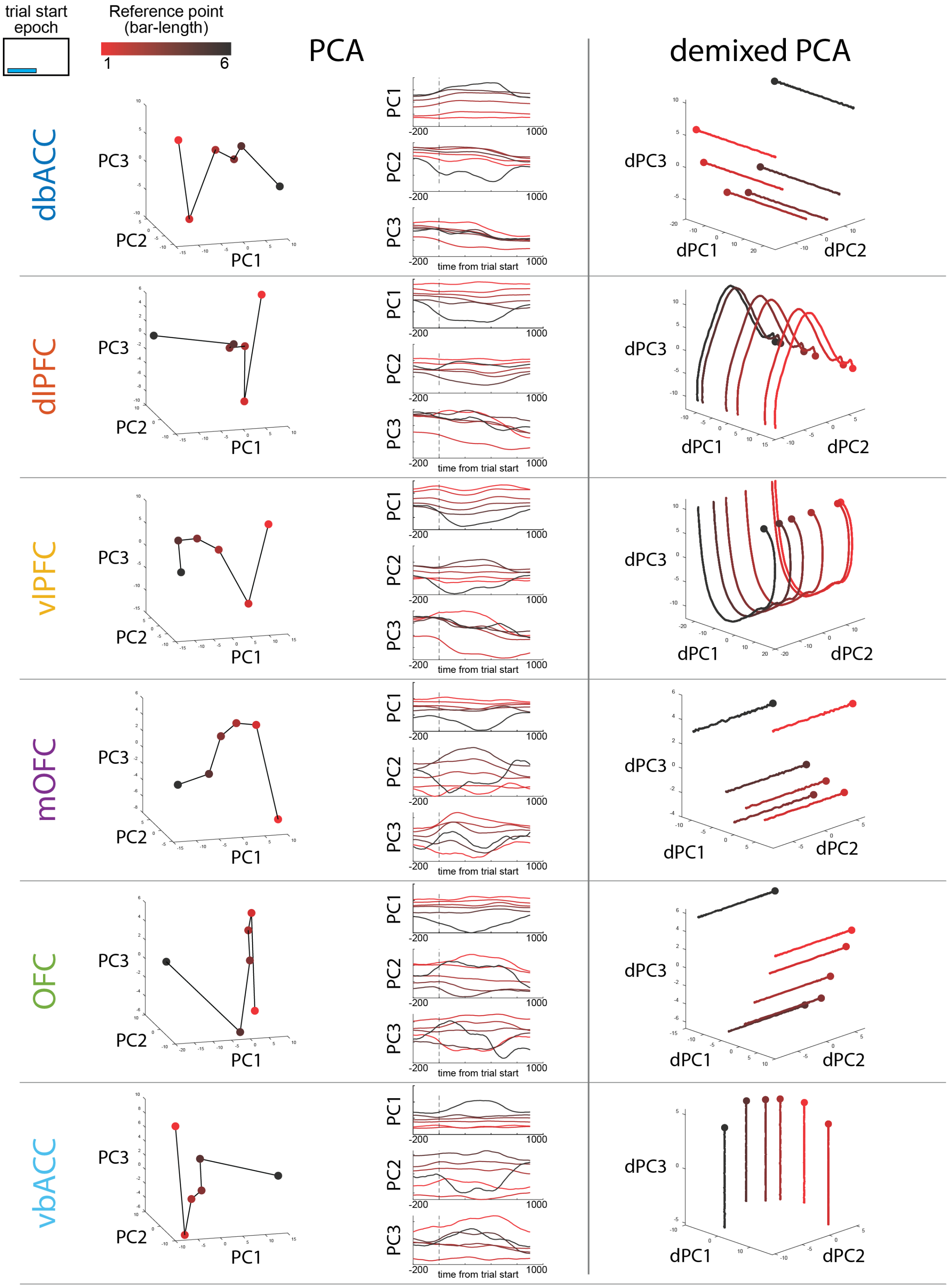

**Figure S1:** Latent dimensional analyses reveal reference-point encoding across brain regions during trial start epoch. Colors correspond to reference levels. (Left column) Factorized population structure, where neural responses reflected reference-point, recovered by PCA. (Middle column) PCA trajectories on the first three principal components. (Right column) dPCA trajectories show the population responses through time across different reference levels. Circle indicates the starting point (200ms before trial start).

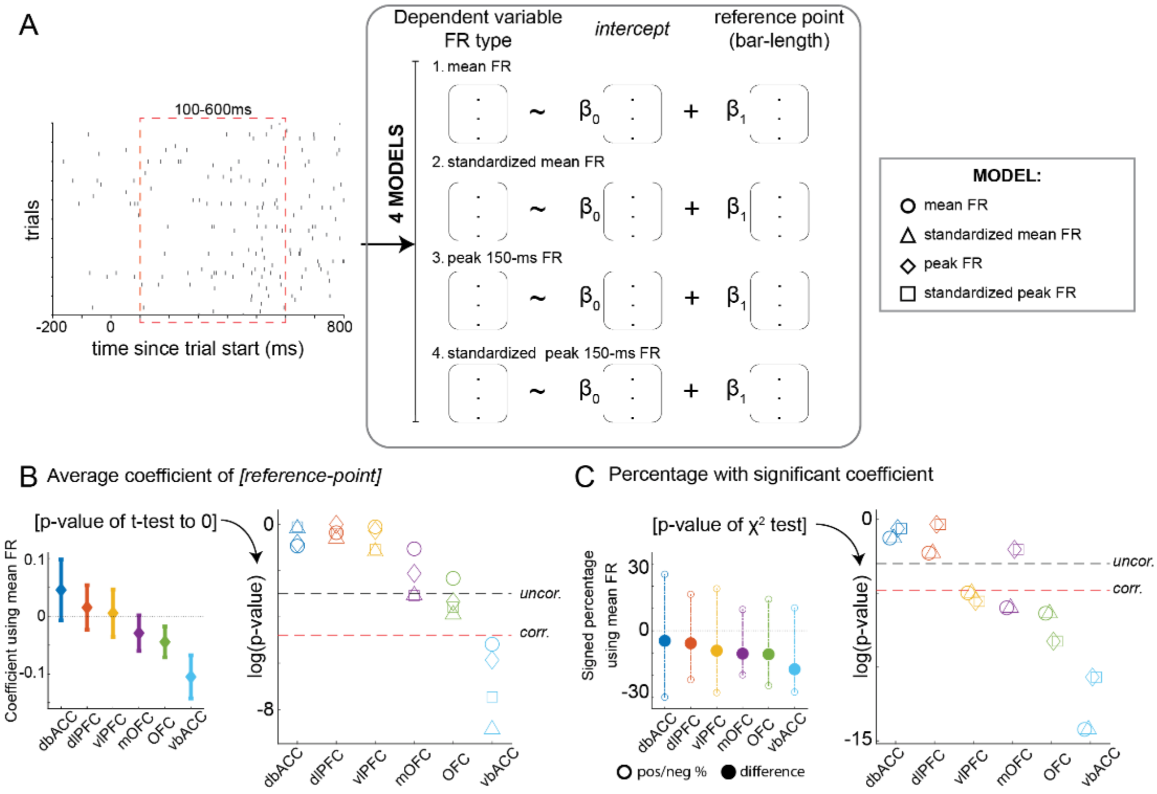

**Figure S2:** The population level reference point signal in vbACC during trial start epoch is robust regardless of FR calculation.

A. Analysis plan. (Left) Time window for analysis. (Middle) For each neuron, we calculated firing-rate (FR) in 4 different ways: (1) average spike count as shown in figure 3 (mean FR); (2) standardized mean FR across trials (standardized mean FR); (3) calculated peak FR of 150ms window (peak FR); (4) standardized peak FR across trials (standardized peak FR). Each of these types of FR were used to fit regression models.
B. (Left) Mean ± SEM for coefficient values of reference point, adapted from figure 3C. For each model, we tested the average coefficient values of all neurons of each brain area against 0 using a one-sample t-test. (Right) P-values of the t-test for each model across brain areas. The black dashed line indicates the uncorrected significance threshold ($p = 0.05$); the red dashed line represents the Bonferroni-corrected threshold for multiple comparisons across six brain areas ($p = 0.0083$). Similar for other figures.
C. (Left) The percentage of population that had significant coefficient value for reference point signal per brain area, adapted from figure 3D. For each model, we tested whether there is a significant difference between positive and negative percentage using a $\chi^2$ test. (Right) P-values of $\chi^2$ test for each model across brain areas.

B-right-panel and C-right-panel are supplements to figures 3C and D.

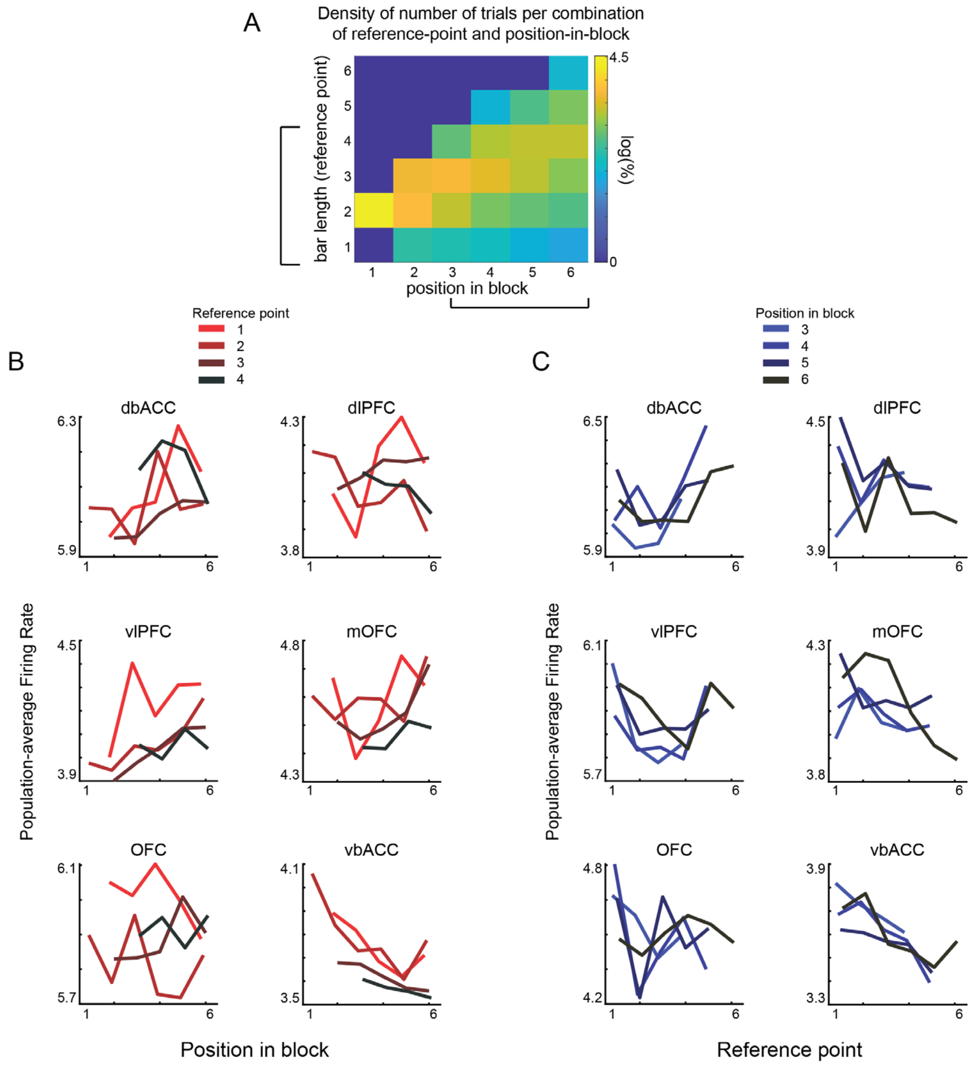

**Figure S3**: Population-average activities clustered by reference points and position in block.

A. Heatmap showing the density of trials for each combination of reference point level and position in block. Based on this distribution, we analyzed trials at reference levels 1–4 (which span a broad range of positions in block; panel B) and trials from positions 3–6 (which span a broad range of reference levels; panel C).
B. Population-average activity across different positions in the block, grouped by reference point level. vbACC shows some sensitivity to position in block, but also clearly reflects a graded response to reference point.
C. Population-average activities across reference point levels grouped by different position in block. vbACC activities responded to reference point but there's no difference between position in blocks.

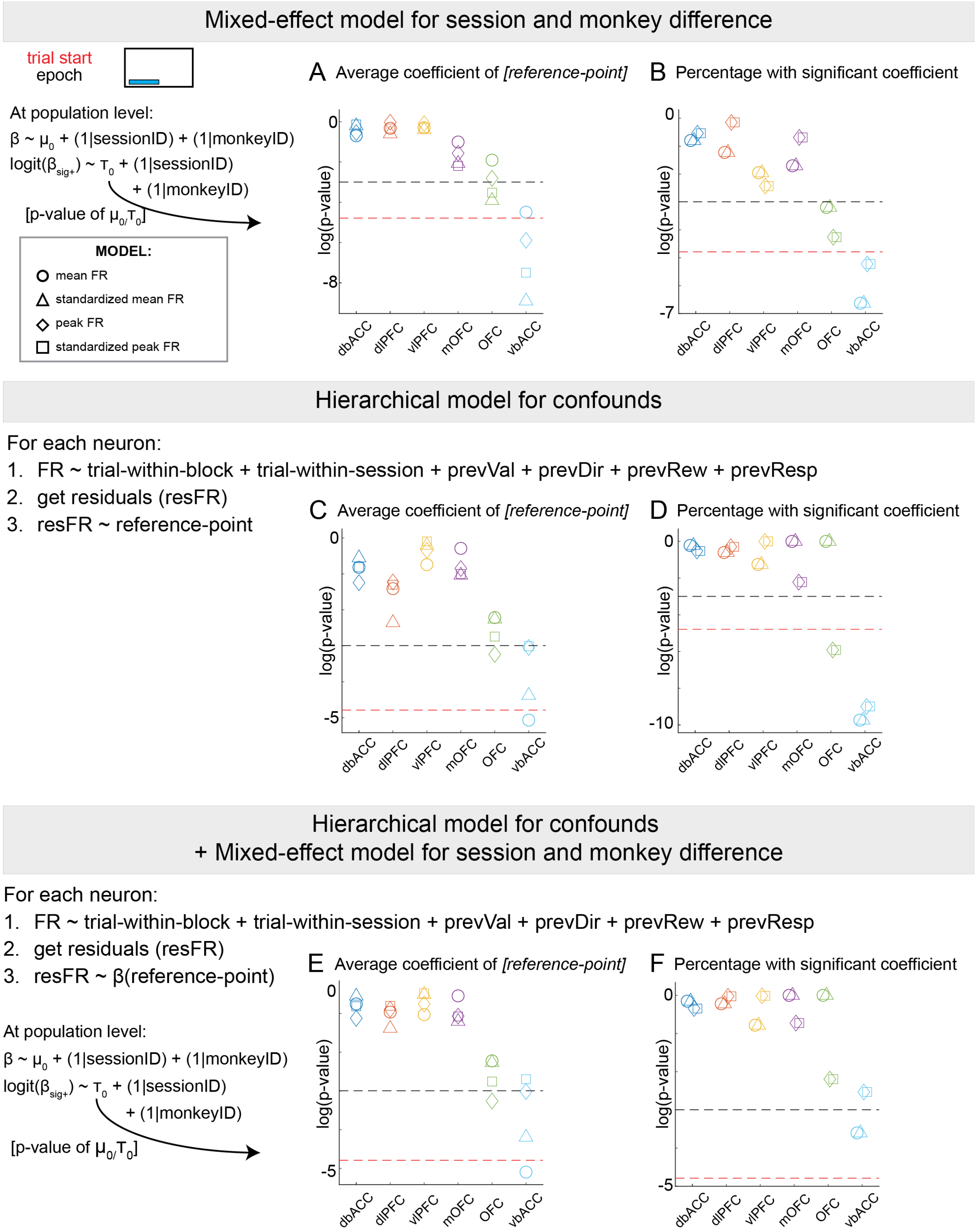

**Figure S4**: The population level reference point signal in the vbACC remains robust even after controlling for potential confounding variables.

For panels A and B, we used a mixed-effects model to account for subject identity when analyzing reference point coefficients across neurons within each brain region. This model tested two hypotheses: (1) whether the population-average coefficient ($\mu_0$) significantly differs from zero, and (2) whether, among neurons with significant coefficients, there is a directional bias ($\tau_0 \neq 0$).

A. P-value of $\mu_0$ for each model across brain areas. The black dashed line indicates the uncorrected significance threshold (p = 0.05); the red dashed line represents the Bonferroni-corrected threshold for multiple comparisons across six brain areas (p = 0.0083). Similar for other figures.
B. P-values of the $\tau_0$ for each model across brain areas.

For panels C and D, we implemented a hierarchical model to control for potential confounding factors (see Methods for details). For each neuron, we first regressed out the influence of these confounds, then modeled the residual variance using the reference point. This allowed us to test whether the vbACC continues to show homogeneous encoding of the reference point signal after controlling for confounds.

C. P-value of t-test for each model across brain areas for coefficient values of reference point.
D. P-value of $\chi^2$ test for each model across brain areas for bias in percentage of population that had significant coefficient value for reference point.

For panels E and F, we combined both approaches described above. First, we used a hierarchical model to control for confounding factors. We then applied a mixed-effects model to account for subject identity when analyzing reference point coefficients across neurons across brain regions.

E. P-value of $\mu_0$ for each model across brain areas.
F. P-value of $\tau_0$ for each model across brain areas.

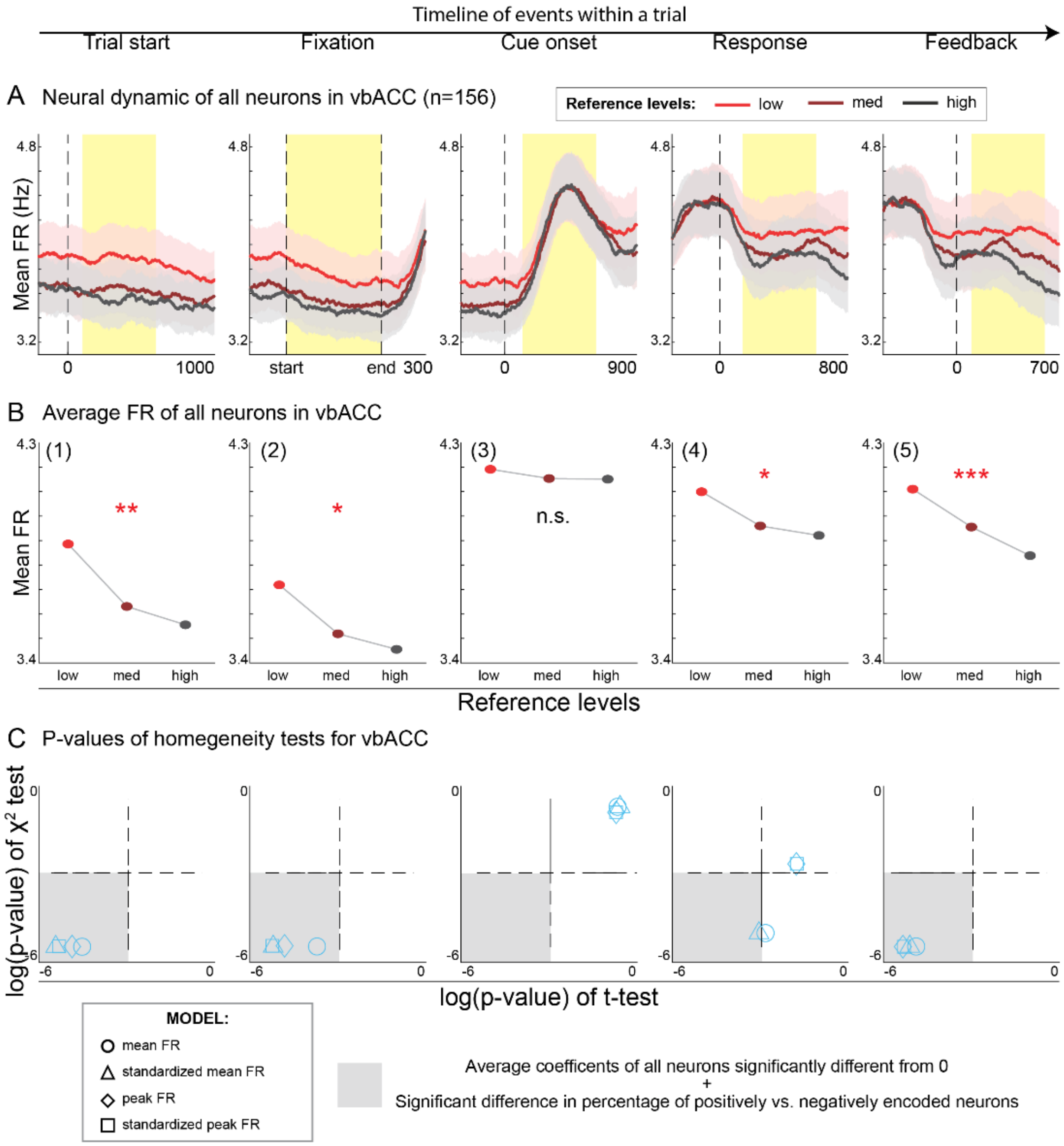

**Figure S5**: A sustained population level reference point signal in the vbACC throughout the trial.

A. (from Fig. 3E) Average firing rate of all neurons in vbACC across all events of a trial. Line indicates mean and shaded area indicates SEM. Yellow shaded area indicates the time window used for analysis.
B. Average firing rate within analysis windows across all events separated based on reference levels. P-values of repeated-measured ANOVA test for each epoch: (1) 0.0027; (2) 0.0114; (3) 0.8614; (4) 0.0210; (5) $9.05 \times 10^{-5}$.
C. P-values of the t-test and $\chi^2$ test for reference point encoding for each model across all epochs. Shaded areas indicated significant homogeneous encoding of reference point. For visualization purpose, here we display (p-value + 0.003) to keep the y-axis bound between [-5.8, 0.003] and roughly equal range for above and below p-value of 0.05.

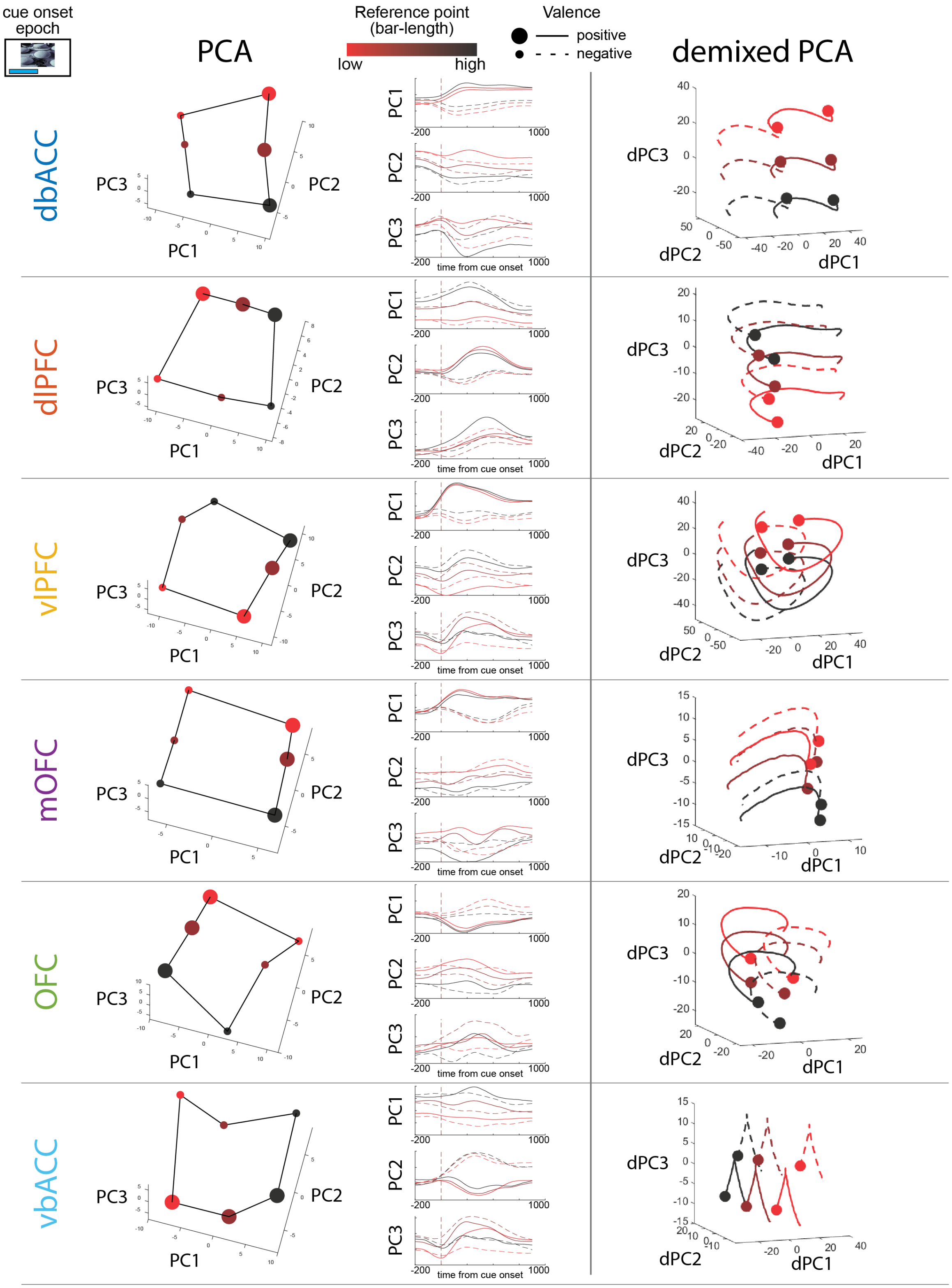

**Figure S6:** Latent dimensional analyses reveal reference-dependent cue-valence encoding across brain regions during cue onset epoch. Colors correspond to reference levels; size and line-style corresponds to cue valence. (Left column) Factorized population structure, where neural responses reflected reference-dependent cue valence signal, recovered by PCA. (Middle column) PCA trajectories on the first three principal components. (Right column) dPCA trajectories show the population responses through time across different reference levels and cue valence. Circle indicates the starting point (200ms before cue onset).

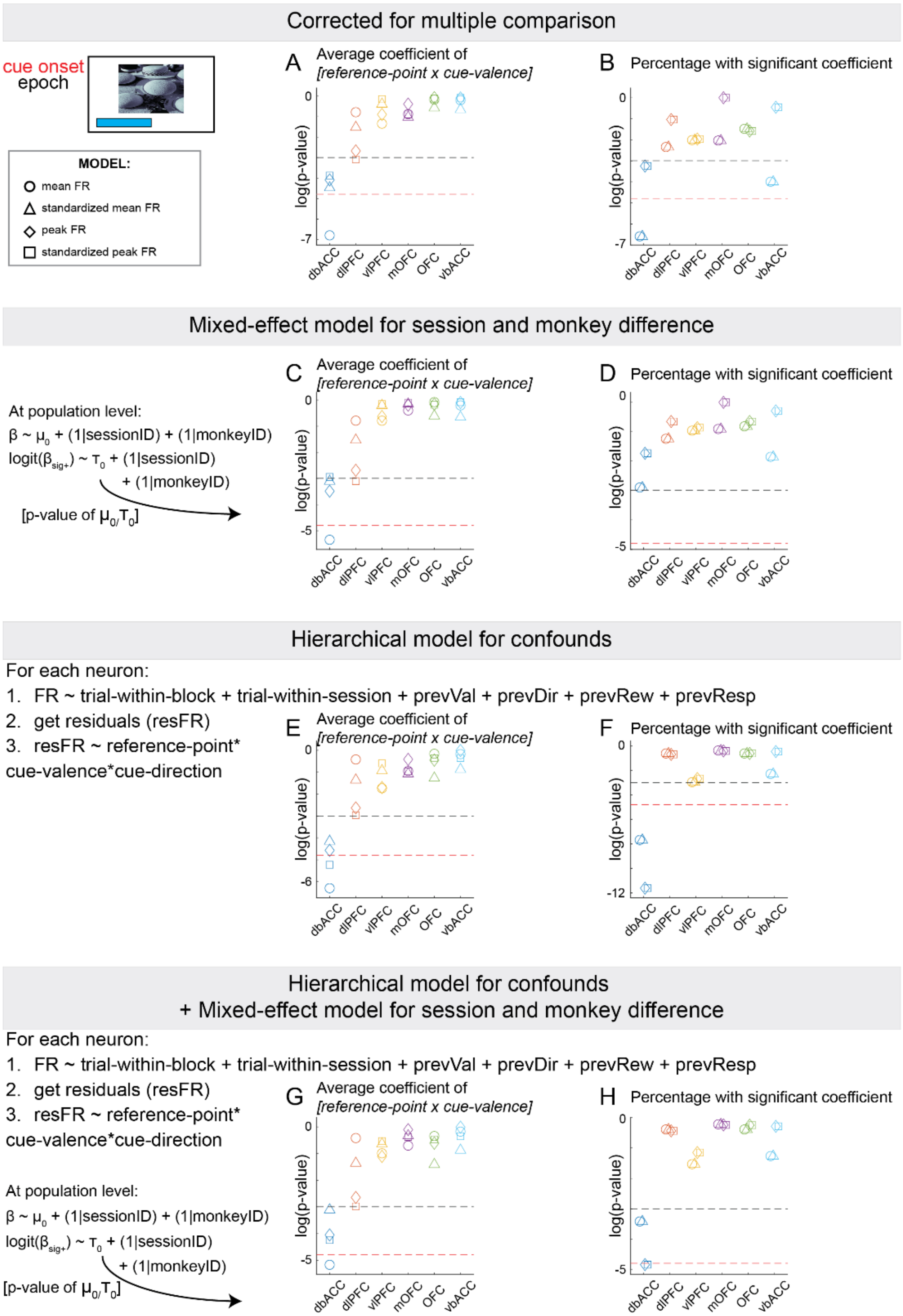

**Figure S7:** The population level reference-dependent cue-valence signal in the dbACC remains robust even after controlling for potential confounding variables.

For panels A and B, we tested the homogenous signal using different measurements of firing rate for the regression models.

A. P-values of the t-test for each model across brain areas for coefficient values of reference-dependent cue-valence signal (interaction term [*reference-point* × *cue-valence*]). The black dashed line indicates the uncorrected significance threshold ($p = 0.05$); the red dashed line represents the Bonferroni-corrected threshold for multiple comparisons across six brain areas ($p = 0.0083$). Similar for other figures.
B. P-values of $\chi^2$ test for each model across brain areas.

For panels C and D, we used a mixed-effects model to account for subject identity.

C. P-value of $\mu_0$ for each model across brain areas.
D. P-values of the $\tau_0$ for each model across brain areas.

For panels E and F, we implemented a hierarchical model to control for potential confounding factors (see Methods for details). For each neuron, we first regressed out the influence of these confounds, then modeled the residual variance using the reference point, cue valence, cue direction and their interaction terms.

E. P-values of the t-test for each model across brain areas for coefficient values of reference-dependent cue-valence signal (interaction term [*reference-point* × *cue-valence*]).
F. P-value of $\chi^2$ test of each model for across brain areas.

For panels G and H, we combined both approaches described above. First, we used a hierarchical model to control for confounding factors. We then applied a mixed-effects model to account for subject identity when analyzing the reference-dependent cue-valence signal.

G. P-value of $\mu_0$ for each model across brain areas.
H. P-value of $\tau_0$ for each model across brain areas.

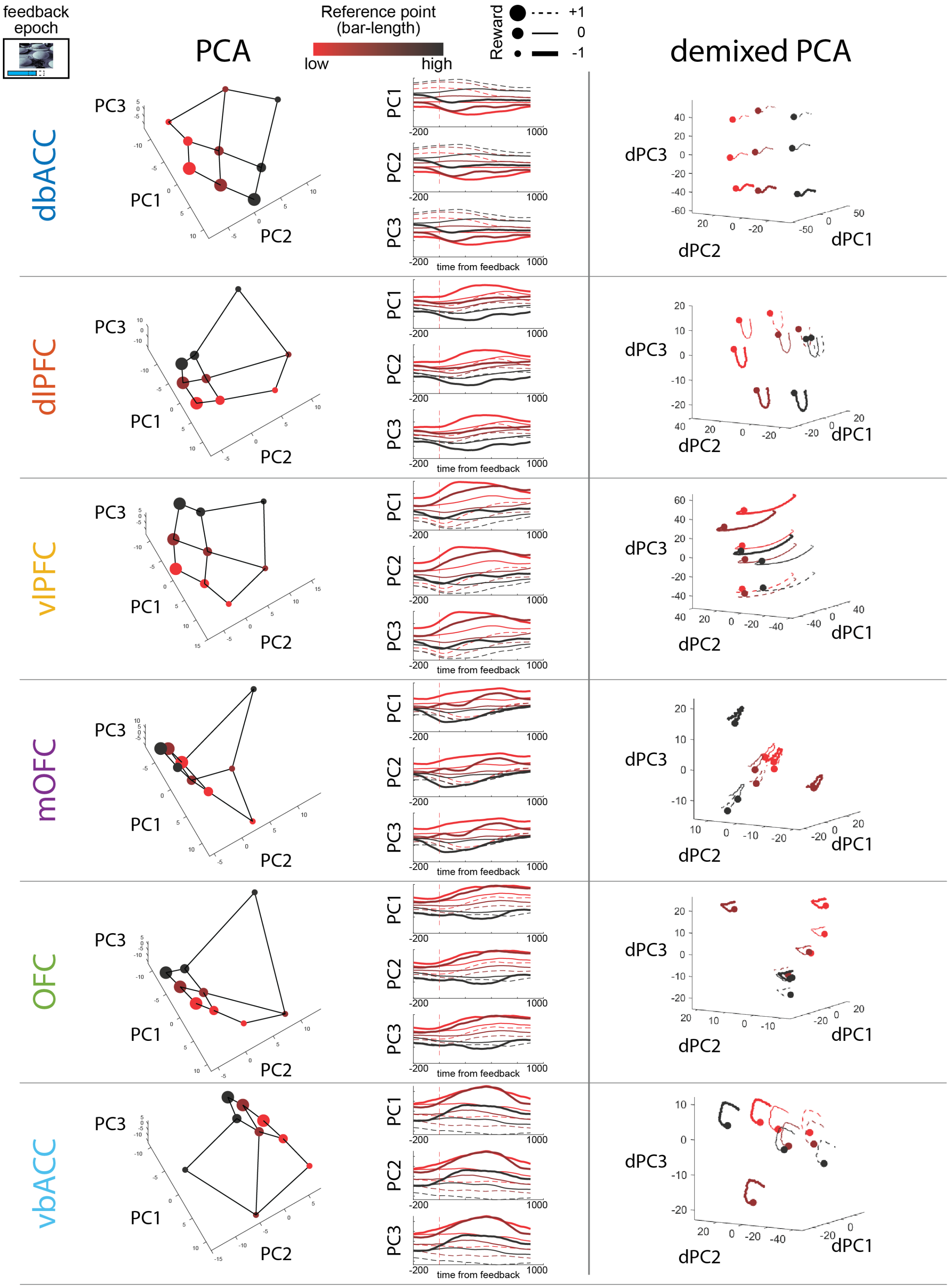

**Figure S8:** Latent dimensional analyses reveal reference-dependent outcome encoding across brain regions during feedback epoch. Colors correspond to reference levels; size and line-style corresponds to outcome. (Left column) Factorized population structure, where neural responses reflected reference-dependent outcome signal, recovered by PCA. (Middle column) PCA trajectories on the first three principal components. (Right column) dPCA trajectories show the population responses through time across different reference and outcome levels. Circle indicates the starting point (200ms before feedback onset).

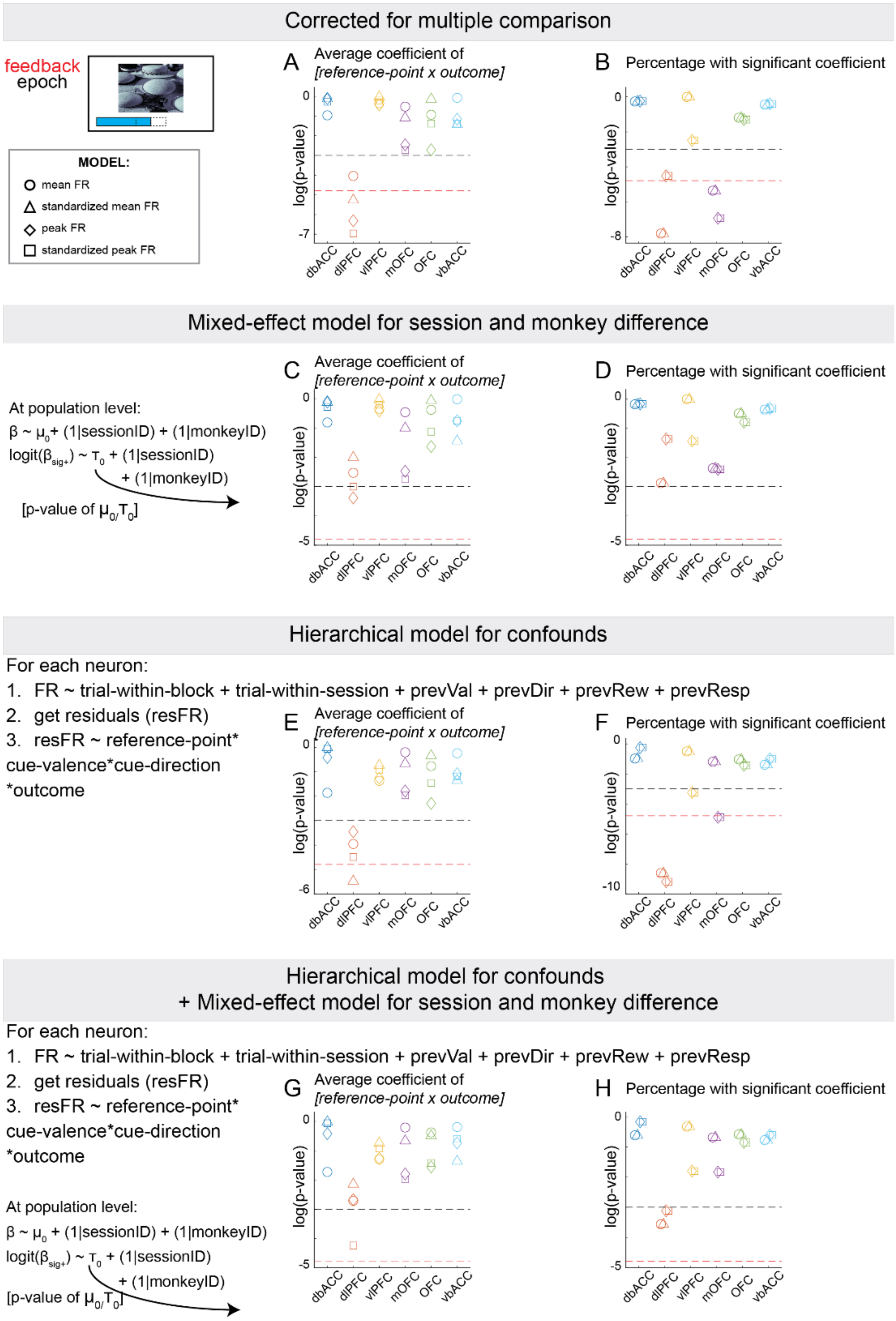

**Figure S9:** The population level reference-dependent outcome signal in the dlPFC remains robust even after controlling for potential confounding variables.

For panels A and B, we tested the homogenous signal using different measurements of firing rate for the regression models.

A. P-values of the t-test for each model across brain areas for coefficient values of reference-dependent cue-valence signal (interaction term [*reference-point* × *outcome*]). The black dashed line indicates the uncorrected significance threshold (p = 0.05); the red dashed line represents the Bonferroni-corrected threshold for multiple comparisons across six brain areas (p = 0.0083). Similar for other figures.
B. P-values of $\chi^2$ test for each model across brain areas.

For panels C and D, we used a mixed-effects model to account for subject identity.

C. P-value of $\mu_0$ for each model across brain areas.
D. P-values of the $\tau_0$ for each model across brain areas.

For panels E and F, we implemented a hierarchical model to control for potential confounding factors (see Methods for details). For each neuron, we first regressed out the influence of these confounds, then modeled the residual variance using the reference point, cue valence, cue direction, outcome and their interaction terms.

E. P-values of the t-test for each model across brain areas for coefficient values of reference-dependent cue-valence signal (interaction term [*reference-point* × *cue-valence*]).
F. P-value of $\chi^2$ test of each model for across brain areas.

For panels G and H, we combined both approaches described above. First, we used a hierarchical model to control for confounding factors. We then applied a mixed-effects model to account for subject identity when analyzing the reference-dependent cue-valence signal.

G. P-value of $\mu_0$ for each model across brain areas.
H. P-value of $\tau_0$ for each model across brain areas.

### Supplemental Note 1

*Introduction*

This study seeks to provide direct evidence for a neural representation of the decisional reference point, a central construct in behavioral economics. In the main text, we defined the accumulated reward (presented as bar length) as the objective reference point, following Kahneman and Tversky's original work (Kahneman and Tversky 1979). However, monkeys may instead rely on a subjective reference point that evolves trial-by-trial and is only partly aligned with accumulated reward. Such latent variables could (1) better capture behavior, (2) more closely match single-neuron activity, and (3) potentially reveal distinct population-level computations across brain regions

To examine this possibility, we evaluated four alternative models of subjective reference points:

**Model 1:** Log-transformed bar length (nonlinear transformation)
**Model 2:** Reference point inferred from reaction time
**Model 3:** Bayesian reinforcement learning (BRL) with *a prior*
**Model 4:** BRL with discounted predictions of future expected reward

Each model was compared to the baseline **Model 0** (accumulated reward as reference point). We assessed them using three criteria: behavioral fit, explanatory power for single-neuron activity, and evidence for population-level encoding.

For each model, we provide:

- *Methods* **-** Construction of the subjective reference point (SRP).
- *Results* **-** Performance in explaining behavior, single-neuron activity, and population-level encoding.
- *Discussion* **-** Relation to the original findings.

All new analyses followed the same regression framework used in the main text, replacing the accumulated reward with the SRP estimated from each model.

1. <u>Non-linear transformation of bar-length</u>

*Methods*

Our experimental design did not allow a direct estimate of the curvature of the utility functions used by Kahneman and Tversky. Instead, we adopted a widely used nonlinear transformation in the literature first proposed by Bernoulli, the log transformation (Bernoulli 1967).

$$\mathrm{SRP} = \log\,(\mathrm{bar} - \mathrm{length})$$

*Results*

a) Relationship with accumulated reward

As expected, the SRP derived from the log transformation was strongly correlated with the objective accumulated reward. This confirms that the transformation preserved much of the structure of the bar-length signal while introducing nonlinearity ($\rho = 0.9693, \mathrm{p} \simeq 0$).

b) Behavioral performance explained

We compared behavioral model fits using either the log-transformed SRP (Model 1) or the accumulated reward (Model 0). Results are summarized in Supplemental Note Table 1. Overall, the log-transformed SRP provided a better fit to both reaction time and percent correct than the objective bar length.

| | REACTION TIME | | PERCENT CORRECT | |
|---|---|---|---|---|
| | Model 0 | Model 1 | Model 0 | Model 1 |

| Coefficients<br>(Mean<br>SE<br>p-value) | | | | |
|---|---|---|---|---|
| intercept | 1002.5<br>108.33<br>$2.27\times10^{-20}$ | 1344.4<br>110.17<br>$3.47\times10^{-34}$ | 1.24<br>0.6128<br>0.0428 | 0.81<br>0.6160<br>0.1886 |
| Reference Point (RP) | -123.87<br>6.05<br>$8.63\times10^{-93}$ | -534.61<br>21.97<br>$8.02\times10^{-130}$ | 0.23<br>0.0197<br>$3.66\times10^{-32}$ | 0.84<br>0.0658<br>$1.66\times10^{-37}$ |
| Cue-valence | -382.99<br>25.10<br>$2.00\times10^{-52}$ | -621.73<br>39.49<br>$1.13\times10^{-55}$ | 1.66<br>0.0973<br>$2.93\times10^{-65}$ | 1.90<br>0.1458<br>$1.23\times10^{-38}$ |
| Cue-valence × RP | 78.685<br>8.20<br>$9.04\times10^{-22}$ | 356.15<br>29.59<br>$2.61\times10^{-33}$ | -0.16<br>0.0335<br>$2.43\times10^{-6}$ | -0.53<br>0.1128<br>$3.17\times10^{-6}$ |
| Model fit | | | | |
| $R^2$ | 0.0524 | 0.0570 | 0.0696 | 0.0703 |

**Supplemental Note Table 1**: Comparison of behavioral performance explained between Model 1 and Model 0.

c) Single neural activity explained

To test whether the log-transformed SRP improved the explanation of single-neuron activity, we repeated the regression analyses from the main text, substituting the SRP for the accumulated reward. For each neuron, we compared the variance explained by Models 0 and 1. Unlike behavior, neural activity was consistently better explained by Model 0. This was true both across all neurons and when restricting the analysis to vbACC neurons (Supplemental Note Fig. 1).

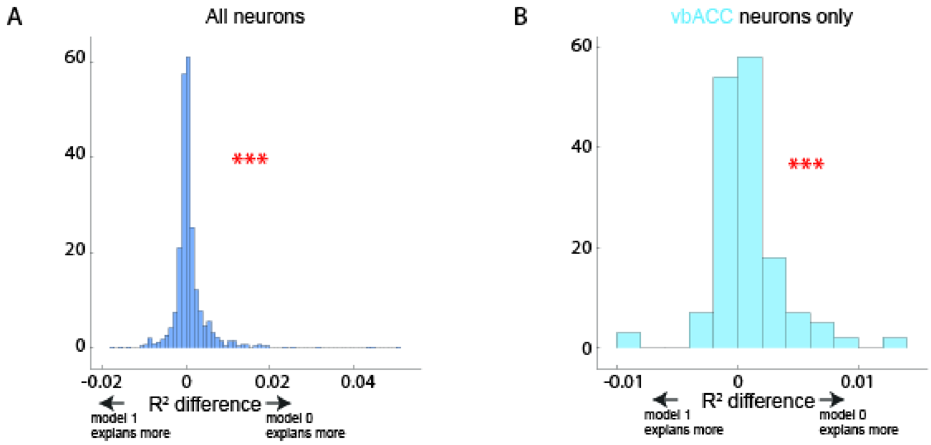

**Supplemental Note Figure 1**: $R^2$ explained between original model (model 0) and SRP using log-transformation (model 1). (A) Comparison between all neurons. (B) Comparison between vbACC, the area we identified the population-level encoding of the reference point, only.

d) Population-average

At the population level, we examined whether Model 1 captured reference-point encoding across brain regions. Consistent with the main analysis, significant population-level encoding was found only in the vbACC (Supplemental Note Fig. 2). Thus, despite differences in behavioral and single-neuron fits, both Models 0 and 1 converge on the same population-level conclusion.

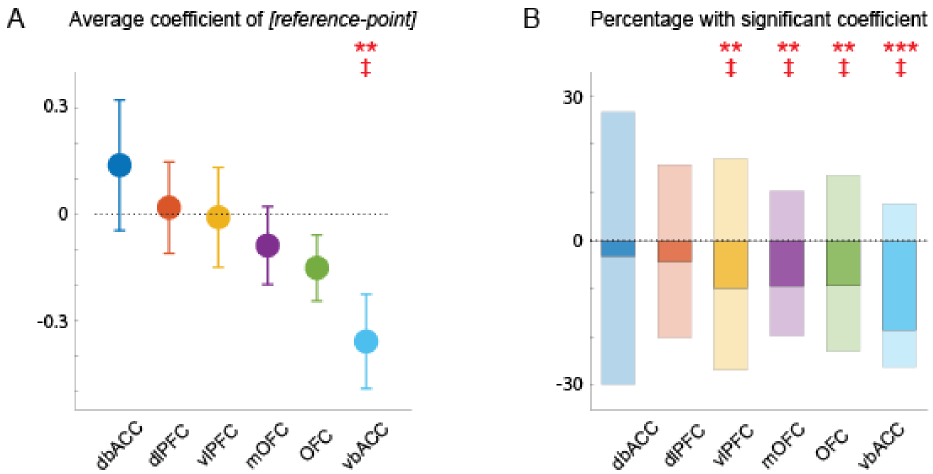

**Supplemental Note Figure 2**: Population analysis using SRP from Model 1. (A) Mean ± SEM for all coefficient values of subjective reference point (*Model 1*) of neurons grouped by brain area. (B) The percentage of population with significantly positive and negative coefficient values for subjective reference point (*Model 1*) per brain area (lighter shade) and the difference between these two numbers (darker shade). Error bars represent mean ± SEM. *p ≤ 0.05, **p ≤ 0.01, ***p ≤0.001, n.s. p > 0.05. ‡p ≤ 0.0083 (Bonferroni-corrected for 6 brain areas).

*Discussion*

In summary, applying a log transformation to bar length improved the fit to behavioral data but did not outperform the objective reference point in explaining single-neuron activity. Importantly, both approaches yielded the same population-level result: robust encoding of the reference point only in vbACC.

2. <u>Subjective reference point inferred from reaction time</u>

*Methods*

Decision making requires not only selecting an action but also determining its vigor (speed), a process theorized to maximize long-term reward rate (Niv, Daw et al. 2007). We adopted a previously proposed framework linking accuracy and response time. We made two assumptions. First, monkeys seek to maximize reward rate over the session. This allows us to write down the equation:

$$V^*(S) = \max_\tau \left[ -\rho - \frac{K}{\tau} + U(R)P(R|S,\tau) - \tau\bar{R} + V^*(S') \right]$$

where $(-\rho - \frac{K}{\tau})$ is the cost of response consisting of a fixed and time-dependent term; $U(R)$ is the utility of potential reward with probability $P(R|S,\tau)$ dependent on the state and response time, the opportunity cost $(-\tau\bar{R})$ depending on the response time and the average reward rate, $\bar{R}$, mathematically equivalent to the reference point. Second, we made assumption about the relationship between performance and response time:

$$P(R|S,\tau) = \frac{1}{1 + \exp(-\beta 0 - \beta 1 * \tau)}$$

Following the first assumption, we can find the maximal expected value by taking the derivative:

$$\frac{dV^*(S)}{d\tau} = \frac{K}{\tau^2} - U(R)\frac{dP(R|S,\tau)}{d\tau} - \bar{R}$$

Setting this to zero and solving for $\bar{R}$ gives the optimal solution:

$$\mathrm{SRP} = \bar{R} = \frac{K}{\tau^2} - U(R)\frac{dP(R|S,\tau)}{d\tau}$$

We estimated the free parameters $\beta 0$ and $\beta 1$ by fitting the $P(R|S,\tau)$ at session level and performed grid search for values of $K$ and $U(R)$ to maximize the correlation between this SRP and the objective bar length.

*Results*

a) Relationship with accumulated reward

Despite parameter tuning, the SRP from this model showed only a weak, though statistically significant, correlation with accumulated reward ($\rho = 0.0209, p \simeq 4.3 \times 10^{-5}$).

b) Behavioral performance explained

Model fits are summarized in Supplemental Table 2. Both models captured reaction time variance, but Model 0 (objective bar length) performed substantially better. Interestingly, Model 2's SRP predicted accuracy in the opposite direction of Model 0: negative coefficients rather than positive.

| | REACTION TIME | | PERCENT CORRECT | |
|---|---|---|---|---|
| | Model 0 | Model 2 | Model 0 | Model 2 |
| Coefficients (Mean SE p-value) | | | | |
| intercept | 1002.5<br>108.33<br>$2.27\times10^{-20}$ | 772.8<br>111.89<br>$5.02\times10^{-12}$ | 1.24<br>0.6128<br>0.0428 | 1.94<br>0.6155<br>0.0016 |
| Reference Point (RP) | -123.87<br>6.05<br>$8.63\times10^{-93}$ | -50.65<br>6.91<br>$2.42\times10^{-13}$ | 0.23<br>0.0197<br>$3.66\times10^{-32}$ | -0.0251<br>0.0210<br>0.2306 |
| Cue-valence | -382.99<br>25.10<br>$2.00\times10^{-52}$ | -276.5<br>24.55<br>$2.28\times10^{-29}$ | 1.66<br>0.0973<br>$2.93\times10^{-65}$ | 1.33<br>0.1024<br>$8.78\times10^{-39}$ |
| Cue-valence × RP | 78.685<br>8.20<br>$9.04\times10^{-22}$ | 49.31<br>9.02<br>$4.58\times10^{-8}$ | -0.16<br>0.0335<br>$2.43\times10^{-6}$ | -0.04<br>0.0367<br>0.2858 |
| Model fit | | | | |
| $R^2$ | 0.0524 | 0.0417 | 0.0696 | 0.0680 |

**Supplemental Note Table 2**: Comparison of behavioral performance explained between Model 2 and Model 0

c) Single neural activity explained

We repeated the neural regression analyses with the SRP from Model 2. Across all neurons, including vbACC, variance explained was markedly lower than with Model 0 (Supplemental Note Fig. 3).

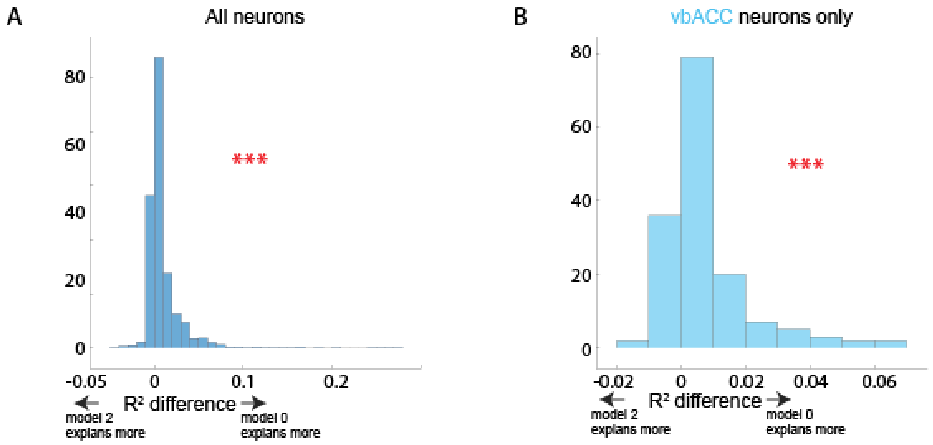

**Supplemental Note Figure 3**: $R^2$ explained between original model (model 0) and SRP inferred from reaction time (model 2). (A) Comparison between all neurons. (B) Comparison between vbACC, the area we identified the population-level encoding of the reference point, only.

d) Population-average

Population-level analysis with Model 2 showed no significant encoding of SRP in any brain region (Supplemental Note Fig. 4).

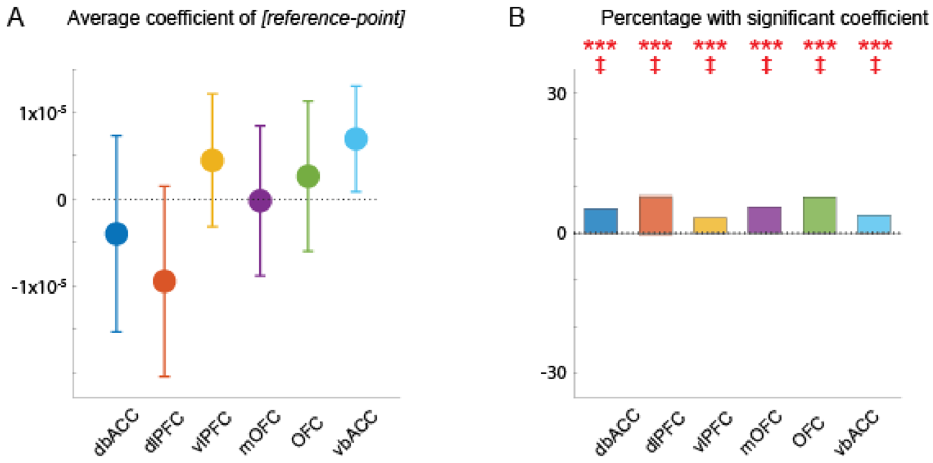

**Supplemental Note Figure 4**: Population analysis using SRP from Model 2. (A) Mean ± SEM for all coefficient values of subjective reference point (*Model 2*) of neurons grouped by brain area. (B) The percentage of population with significantly positive and negative coefficient values for subjective reference point (*Model 2*) per brain area (lighter shade) and the difference between these two numbers (darker shade). Error bars represent mean ± SEM. *$p \leq 0.05$, **$p \leq 0.01$, ***$p \leq 0.001$, n.s. $p > 0.05$. ‡$p \leq 0.0083$ (Bonferroni-corrected for 6 brain areas).

*Discussion*

Although this model offers a principled way to link accuracy and vigor, it relies on restrictive assumptions. Specifically, it assumes an inverse accuracy-speed trade-off and optimal reward-rate maximization, assumptions that do not fully hold in our data. For example, we observed performance drift across session time, suggesting non-optimal behavior. We attempted to account for this by regressing out time-in-session effects before fitting, but other residual violations remained.

Consequently, the SRP inferred from reaction times failed to explain behavior, single-neuron activity, or population-level encoding. This suggests that this model is not a suitable approximation of the monkeys' internal reference point.

3. Bayesian reinforcement learning with *a prior*

*Methods*

We start with a standard reinforcement learning approach:

$$V_{t+1} = V_t + \alpha[R - V_t]$$

where $t$ refers to the position of a trial within a block. By assuming the value of the state/ the reference point follows Gaussian distribution, we define the optimal updating rule for α as in (Wetherill 1961):

$$\alpha = \frac{\sigma_{prior}}{\sigma_{prior} + \sigma_{posterior}} \quad (1)$$

In this model, we assumed the monkeys constructed *a prior* at the beginning of each block based on their past sessions and update this expectation throughout each trial until the final reward is delivered (Supplemental Note Fig. 5). Specifically:

$$SRP_0 = \bar{R}$$

$$SRP_t = SRP_{t-1} + \alpha_t[Bar_t - SRP_{t-1}]$$

where *t* is the trial-number within a block; *SRP* is the subjective-reference-point that gets updated every trial; *Bar* refers to bar-length of at each trial. The learning rate $\alpha_t$ is updated based on Equation (1); in which, $\sigma_{prior}$ $(\bar{\sigma})$ is calculated based on the variance of the magnitude of past rewards; and $\sigma_{posterior}$ is calculated at each trial based on the task-structure:

$$\sigma_{posterior} = \sum P(cue - valence)P(correct)R$$

Where $P(cue - valence)$ is the probability of encountering positive or negative cue; $P(correct)$ is the probability of responding correctly to the visual stimulus, calculated monkey level; and *R* is the reward outcome (-1, 0, +1). Let *p* denotes $P(correct)$, which is calculated separately for each monkey, we can derive:

$$\sigma_{posterior} = 0.25 + p(1 - p)$$

Based on the independence between trials in a block, the variance is also dependent. Hence, we can calculate the variance at each trial *t* by summing the variance of numbers of trials left from the current one, i.e.:

$$\sigma_{posterior,\ t} = (7 - t)[0.25 + p(1 - p)]$$

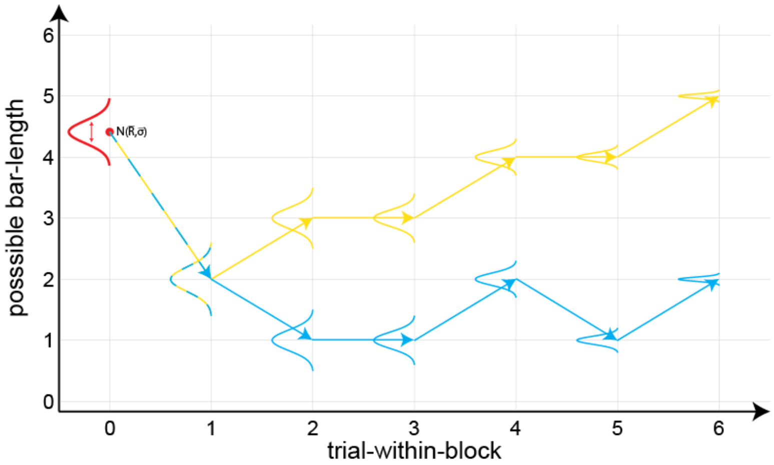

**Supplemental Note Figure 5**: Example of two blocks' trajectories. At the beginning of the block ("trial 0"), monkeys form *a prior* based on past reward deliveries on how much they can expect to get be the end of each block. This *a prior* is updated every trial based on the bar-length that appears.

*Results*

a) Relationship with accumulated reward

The BRL-estimated SRP correlated strongly with accumulated reward, confirming internal consistency ($\rho = 0.9350, p \simeq 0$).

b) Behavioral performance explained

As summarized in Supplemental Note Table 3, the BRL SRP explained behavioral performance comparably well but slightly less effectively than the objective bar length.

| | REACTION TIME | | PERCENT CORRECT | |
|---|---|---|---|---|
| | Model 0 | Model 3 | Model 0 | Model 3 |
| Coefficients (Mean SE p-value) | | | | |
| intercept | 1002.5<br>108.33<br>$2.27\times10^{-20}$ | 1009.8<br>110.03<br>$4.60\times10^{-20}$ | 1.24<br>0.6128<br>0.0428 | 1.23<br>0.6145<br>0.0444 |
| Reference Point (RP) | -123.87<br>6.05<br>$8.63\times10^{-93}$ | -120.22<br>7.71<br>$1.19\times10^{-54}$ | 0.23<br>0.0197<br>$3.66\times10^{-32}$ | 0.22<br>0.0245<br>$2.82\times10^{-19}$ |
| Cue-valence | -382.99<br>25.10<br>$2.00\times10^{-52}$ | -402.04<br>32.18<br>$9.41\times10^{-36}$ | 1.66<br>0.0973<br>$2.93\times10^{-65}$ | 1.73<br>0.1242<br>$3.93\times10^{-44}$ |
| Cue-valence × RP | 78.685<br>8.20<br>$9.04\times10^{-22}$ | 81.20<br>10.32<br>$3.68\times10^{-15}$ | -0.16<br>0.0335<br>$2.43\times10^{-6}$ | -0.17<br>0.0411<br>$2.94\times10^{-5}$ |
| Model fit | | | | |
| $R^2$ | 0.0524 | 0.0473 | 0.0696 | 0.0687 |

**Supplemental Note Table 3**: Comparison of behavioral performance explained between Model 3 and Model 0

c) Single neural activity explained

Across neurons, Model 0 consistently explained more variance in firing than Model 3, including in vbACC (Supplemental Note Fig. 6).

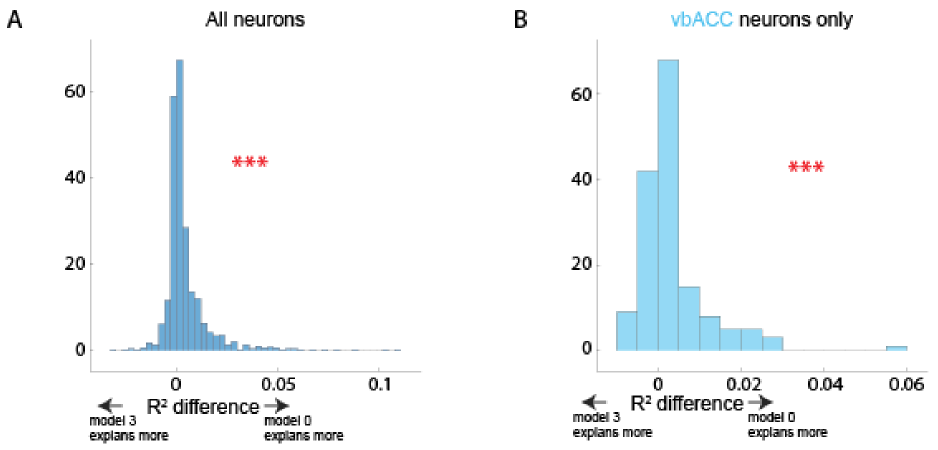

**Supplemental Note Figure 6**: $R^2$ explained between original model (model 0) and SRP inferred based on Bayesian RL with *a prior* (model 3). (A) Comparison between all neurons. (B) Comparison between vbACC, the area we identified the population-level encoding of the reference point, only.

d) Population-average

Population analysis with Model 3 revealed the same result as Model 0: robust SRP encoding in vbACC only (Supplemental Note Fig. 7).

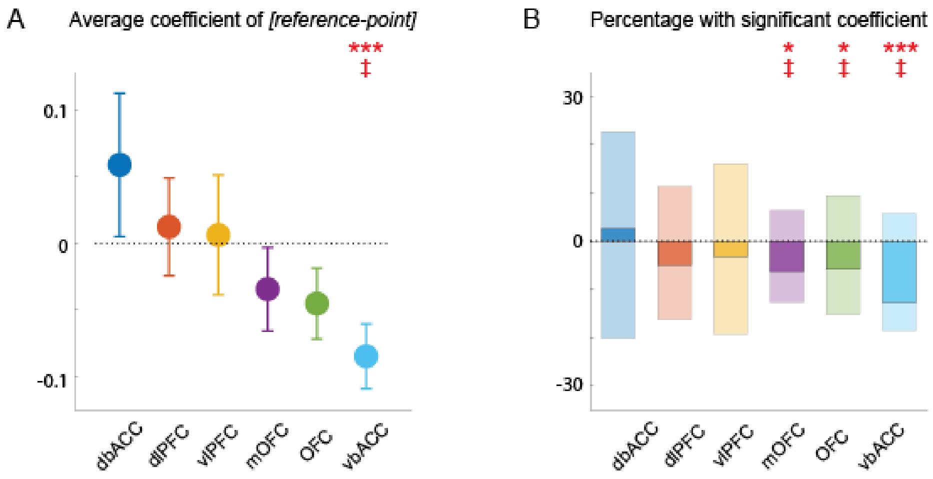

**Supplemental Note Figure 7**: Population analysis using SRP from Model 3. (A) Mean ± SEM for all coefficient values of subjective reference point (*Model 3*) of neurons grouped by brain area. (B) The percentage of population with significantly positive and negative coefficient values for subjective reference point (*Model 3*) per brain area (lighter shade) and the difference between these two numbers (darker shade). Error bars represent mean ± SEM. *p ≤ 0.05, **p ≤ 0.01, ***p ≤0.001, n.s. p > 0.05. ‡p ≤ 0.0083 (Bonferroni-corrected for 6 brain areas).

*Discussion*

The BRL model provided a principled framework for estimating trial-level SRPs based on a prior and ongoing evidence. While it maintained a strong correlation with objective bar length, it did not outperform the simpler model in explaining behavior or neural firing. However, more importantly, both models gave the same conclusion on the population-level encoding of the reference point only in the vbACC.

4. <u>Bayesian reinforcement learning with discounted future expected reward</u>

*Methods*

We examined another standard reinforcement learning approach, focusing on discounting future expected reward:

$$V_{t+1} = V_t + \alpha[\gamma FR_t - V_t]$$

where *t* refers to the position of a trial within a block; and *FR* refers to the final amount of reward predicted by the monkeys (Supplemental Note Fig. 8). Again, by assuming the value of the state/ the reference point follows Gaussian distribution:

$$\alpha = \frac{\sigma_{prior}}{\sigma_{prior} + \sigma_{posterior}}$$

To estimate the future final amount of reward, the predicted amount of increased earned reward per trial is calculated based on the task-structure:

$$\mu = p - 0.5$$

$$\sigma = 0.25 + p(1 - p)$$

where *p* denotes the $P(correct)$. Based on the independence between trials in a block, we can calculate the estimated final amount of reward at each trial *t*:

$$\mathrm{FR_t} \sim N(\mu_t, \sigma_t)$$
$$\mu_t = (7 - t)[p - 0.5] \quad (2)$$
$$\sigma_t = (7 - t)[0.25 + p(1 - p)]$$

To summarize, the subjective-reference-point is calculated as:

$$\mathrm{SRP_t} = \mathrm{Bar_t} + \alpha_t[\gamma^{7-t}\mathrm{FR_t} - \mathrm{Bar_t}]$$

where *t* is the trial-number within a block; *SRP* is the subjective-reference-point that gets updated every trial; *Bar* refers to bar-length of at each trial; $\gamma$ is the discounting factor at each trial and is set to 0.9.

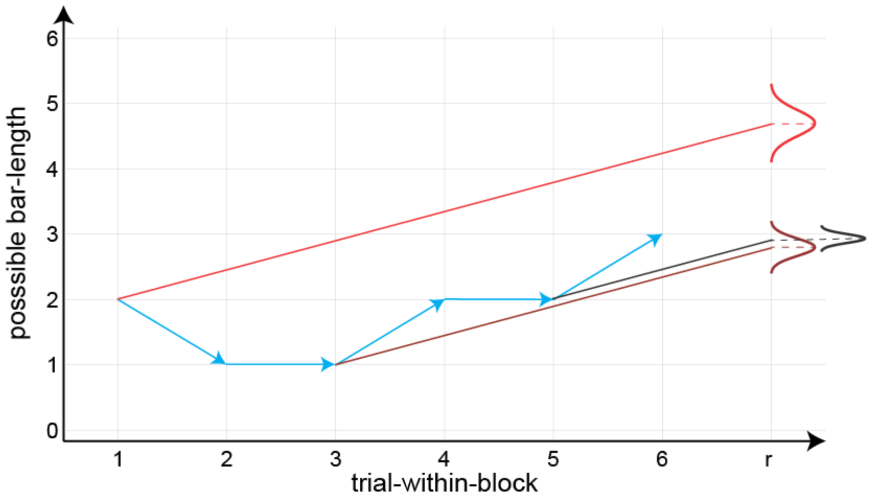

**Supplemental Note Figure 8**: Example of a block's trajectory. At each position within a block, the monkeys can simulate and predict the final reward that they might get. The slope of the predicted increase depends on the monkeys' performance and number of trials left. The stochasticity of the predicted final amount depends on number of trials left. (Equation (2))

*Results*

a) Relationship with accumulated reward

We found a strong correlation between this subjective-reference-point and the accumulated amount of reward ($\rho = 0.9985, p \simeq 0$).

b) Behavioral performance explained

Model 0 explained behavior slightly better than Model 4, though both performed comparably (Supplemental Note Table 4).

| | REACTION TIME | | PERCENT CORRECT | |
|---|---|---|---|---|
| | Model 0 | Model 4 | Model 0 | Model 4 |
| Coefficients (Mean SE p-value) | | | | |
| intercept | 1002.5<br>108.33<br>$2.27\times10^{-20}$ | 1042.4<br>108.85<br>$1.05\times10^{-21}$ | 1.24<br>0.6128<br>0.0428 | 1.15<br>0.6137<br>0.0601 |
| Reference Point (RP) | -123.87<br>6.05<br>$8.63\times10^{-93}$ | -139.44<br>6.91<br>$5.24\times10^{-90}$ | 0.23<br>0.0197<br>$3.66\times10^{-32}$ | 0.27<br>0.0227<br>$1.39\times10^{-31}$ |

| | | | | |
|---|---|---|---|---|
| Cue-valence | -382.99<br>25.10<br>$2.00\times10^{-52}$ | -407.1<br>27.97<br>$7.36\times10^{-48}$ | 1.66<br>0.0973<br>$2.93\times10^{-65}$ | 1.72<br>0.1097<br>$2.82\times10^{-55}$ |
| Cue-valence × RP | 78.685<br>8.20<br>$9.04\times10^{-22}$ | 88.21<br>9.38<br>$5.48\times10^{-21}$ | -0.16<br>0.0335<br>$2.43\times10^{-6}$ | -0.18<br>0.0386<br>$2.98\times10^{-6}$ |
| Model fit | | | | |
| $R^2$ | 0.0524 | 0.0521 | 0.0696 | 0.0696 |

**Supplemental Note Table 4**: Comparison of behavioral performance explained between Model 4 and Model 0

c) Single neural activity explained

In contrast, Model 4 explained significantly more variance in firing rates than Model 0, both across all neurons and within vbACC (Supplemental Note Fig. 9).

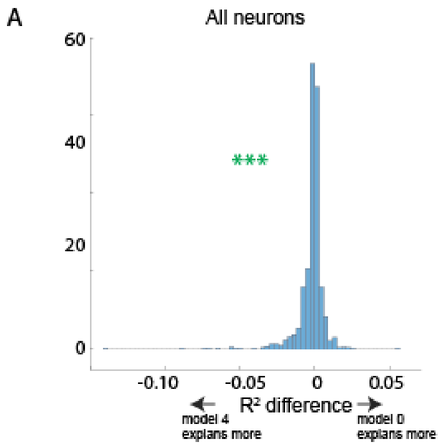

**Supplemental Note Figure 9**: $R^2$ explained between original model (model 0) and SRP inferred based on Bayesian RL with discounted future reward (model 4). (A) Comparison between all neurons. (B) Comparison between vbACC, the area we identified the population-level encoding of the reference point, only.

d) Population-average

Population-level analysis revealed the same conclusion as other models: SRP encoding was present only in the vbACC (Supplemental Note Fig. 10).

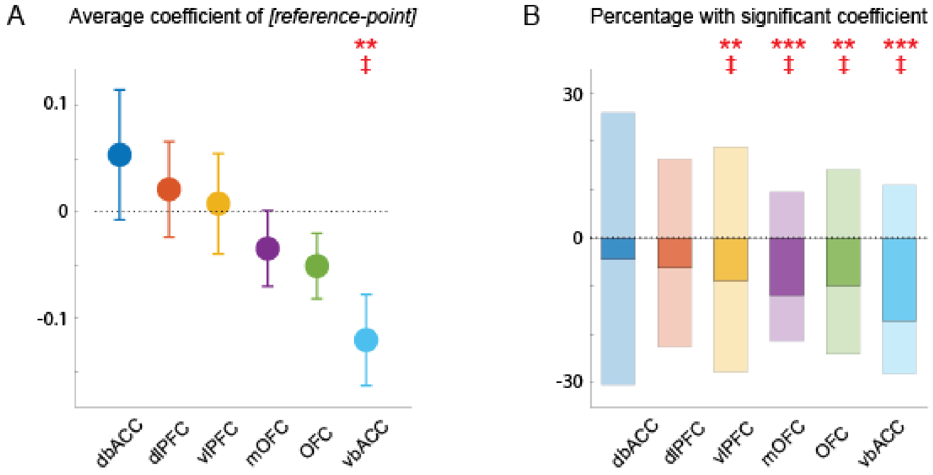

**Supplemental Note Figure 10**: Population analysis using SRP from Model 4. (A) Mean ± SEM for all coefficient values of subjective reference point (*Model 4*) of neurons grouped by brain area. (B) The percentage of population with significantly positive and negative coefficient values for subjective reference point (*Model 4*) per brain area (lighter shade) and the difference between these two numbers (darker shade). Error bars represent mean ± SEM. *$p \leq 0.05$, **$p \leq 0.01$, ***$p \leq 0.001$, n.s. $p > 0.05$. ‡$p \leq 0.0083$ (Bonferroni-corrected for 6 brain areas).

*Discussion*

This model assumes that monkeys form predictions about final block reward and discount them over time. Such an SRP accounted for a greater fraction of neural variance, especially in vbACC, than the objective bar length. However, it did not better explain behavior, raising questions about whether monkeys explicitly rely on this computation for behavior.

Nevertheless, like the other models, it yielded the same population-level conclusion: the vbACC uniquely encodes the reference point.

*General conclusions*

Across four alternative models of subjective reference points, we found differing abilities to explain behavior and single neural activity. Importantly, most models converged on the same key finding: population-level encoding of the reference point was consistently localized to the vbACC. This convergence strongly supports the robustness of our main conclusion.
